# Skin permeable polymer for noninvasive transdermal insulin delivery

**DOI:** 10.1101/2023.05.05.539551

**Authors:** Qiuyu Wei, Zhi He, Jiajia Xiang, Ying Piao, Jianxiang Huang, Yu Geng, Haoru Zhu, Zifan Li, Jiaheng Zeng, Yan Zhang, Hongrui Lu, Quan Zhou, Shiqun Shao, Jianbin Tang, Zhuxian Zhou, Ruhong Zhou, Youqing Shen

## Abstract

Subcutaneous injection of insulin is the current standard medication for many diabetic patients. Convenient and painless noninvasive transdermal insulin delivery has long been pursued but yet succeeded due to no such technologies for large biomacromolecules. We find a tertiary amine oxide-based polyzwitterion, OPDMA, that can efficiently penetrate both the stratum corneum (SC) and viable epidermis into circulation. So its conjugate with insulin, OPDMA-I, applied on the skin can exhibit hypoglycemic effects as efficiently as subcutaneously injected insulin in type-1 diabetic mice and minipigs. The unique pH-dependent cationic-to-zwitterionic transition of OPDMA in the characteristic acidic-to-neutral pH gradient from the skin surface to deep SC enables fast transdermal delivery of OPDMA and its conjugate. On the skin, OPDMA binds to carboxylic acids in the acidic sebum layer, enriching OPDMA-I on the SC. As pH increases in deeper SC layers, binding between OPDMA-I and the skin weakens gradually, allowing for diffusion through inter-corneocyte gaps and penetration into viable epidermis and finally entering the systemic circulation via dermal lymphatic vessels. This process does not alter SC microstructures or cause any physiological changes in the skin. This study represents a groundbreaking example of noninvasive transdermal protein delivery.

## Introduction

Over 425 million patients worldwide with type-1 or advanced type-2 diabetes require exogenous insulin commonly administered *via* intradermal or subdermal injection with syringes, smart insulin pens, insulin pumps, or artificial pancreas^[1]^. However, frequent injections cause pain, needle phobia, and skin complications, thus poor patient compliance^[2]^. Therefore, convenient and noninvasive routes such as oral^[3]^, pulmonary^[4]^, and nasal administrations^[4, 5]^ have been extensively explored but generally suffer low bioavailability, limited dosing flexibility, and safety concerns^[6]^. For instance, inhaled insulin was the first approved as a noninvasive route to deliver insulin but has since been withdrawn from the market.

Transdermal insulin delivery through topical application^[7]^ is advantageous in terms of convenience, high patient compliance, avoiding insulin denature, and minimal first-pass effect^[8, 9]^. However, it is an extremely challenging task due to the formidable barriers presented by the stratum corneum (SC), a 10-20 μm thick matrix composed of dehydrated and dead corneocytes embedded in highly ordered lipid layers^[10]^, as well as the tight junctions in the viable epidermis ^[11]^. The skin barriers selectively allow only small nonpolar molecules with molecular weights less than 500 Da and appropriate hydrophobicities to permeate ^[12]^ while preventing the passage of large biomacromolecules such as insulin ^[13]^. Various techniques have been developed to enhance the skin permeability of insulin^[14]^, including chemical penetration enhancers that disrupt the skin barrier, electrical devices that facilitate penetration^[15]^, ultrasound and jet injection that create transient channels on the skin surface^[16]^, and microneedles that pierce through the stratum corneum into dermal tissue to deposit insulin^[17–19]^. However, these invasive techniques compromise skin integrity and raise infection concerns.

In this study, we demonstrated the highly efficient and noninvasive skin permeation of a polyzwitterion, poly[2-(N-oxide-N,N-dimethylamino)ethyl methacrylate] (OPDMA), and its insulin conjugate (OPDMA-I). Transdermally applied OPDMA-I quickly permeated the skin into the systemic circulation and produced excellent hypoglycemic effects in type-1 diabetic mice and minipigs. The mechanism studies elucidated that OPDMA binds to the acidic sebum layer, leading to an enrichment of OPDMA-I on the SC. Subsequently, it rapidly diffused through the inter-corneocyte lipid lamella, traversed the epidermis and dermis by “hopping” along cell membranes, and finally entered the systemic circulation via dermal lymphatic vessels (Scheme 1).

## Results

### Preparation and characterizations of OPDMA-I

OPDMA was conjugated to insulin, as shown in Fig. S1. OPDMA with a terminal primary amine group (OPDMA-NH_2_) was synthesized according to our previous report, except for using an initiator with a protected amine group^[20]^. The molecular weight was controlled to be 5 kDa. The primary amine was reacted with *N*-(β-maleimidopropyloxy) succinimide ester (BMPS) to introduce a maleimido-group (OPDMA-MAL). Recombinant human insulin was treated with 2-iminothiolane (Traut’s reagent) to introduce a thiol group from the amine group of the lysine residue (insulin-SH). Insulin-SH was coupled with OPDMA-MAL to yield the OPDMA-I conjugate. The PEGylated insulin (PEG-I, PEG’s molecular weight of 5 kDa) was synthesized using the same method^[21]^. The OPDMA-I from the above procedure was confirmed by RP-HPLC (Fig. S2A), MALDI-TOF MS (Fig. S2B), and GPC (Fig. S2C). The conjugation of OPDMA or PEG did not affect the insulin’s secondary structure as characterized by circular dichroism (CD) (Fig. S2D) and the hypoglycemic effect (Fig. S2E).

### Glycemic regulation performance of transdermal OPDMA-I

The *in vivo* glycemic regulation activity of OPDMA-I was assessed in a streptozotocin (STZ)-induced type-1 diabetic mouse model. OPDMA-I, PEG-I, or native insulin solution (0.2 mL, insulin-equivalent concentration: 0.5 mg/mL) in a diffusion cell was applied onto the dorsal skin (1.77 cm^2^) of randomly-grouped diabetic mice (Fig. 1A). The blood glucose levels (BGLs) of the mice were monitored over time. Subcutaneous (*s.c.*) insulin injection sharply lowered the BGLs of diabetic mice from over 400 mg/dL to about 100 mg/dL within 1 h. However, the normoglycemic state could not be maintained, and the mice quickly returned to hyperglycemia after 4 h. Notably, the BGLs of mice exposed to transdermal administration of OPDMA-I dropped below 200 mg/dL (normoglycemic levels) within 1 h, almost as quickly as the *s.c* insulin-treated group, and maintained in a normoglycemic state for up to 12 h with no hypoglycemic events (BGLs below 50 mg/dL). In contrast, transdermally-treated native insulin or PEG-I failed in glycemic regulation (Fig.1B, Fig. S3A). Quantitative analysis of the blood insulin levels by an enzyme-linked immunosorbent assay (ELISA) showed that OPDMA-I rapidly got access to the blood circulatory system upon transdermal application, reaching a maximum of 230 μU/mL at 1 h post-dosing and then declined gradually to 30 μU/mL after 12 h, while few blood insulin was detected in insulin or PEG-I-treated groups (Fig.1C). The relative bioavailability of transdermal OPDMA-I, PEG-I and insulin was determined to be 6.07%,1.35%, and 1.31% compared with *s.c* injection of insulin. The efficient glycemic regulation of OPDMA-I was also validated in healthy mice (Fig. 1D).

**Fig.1.**
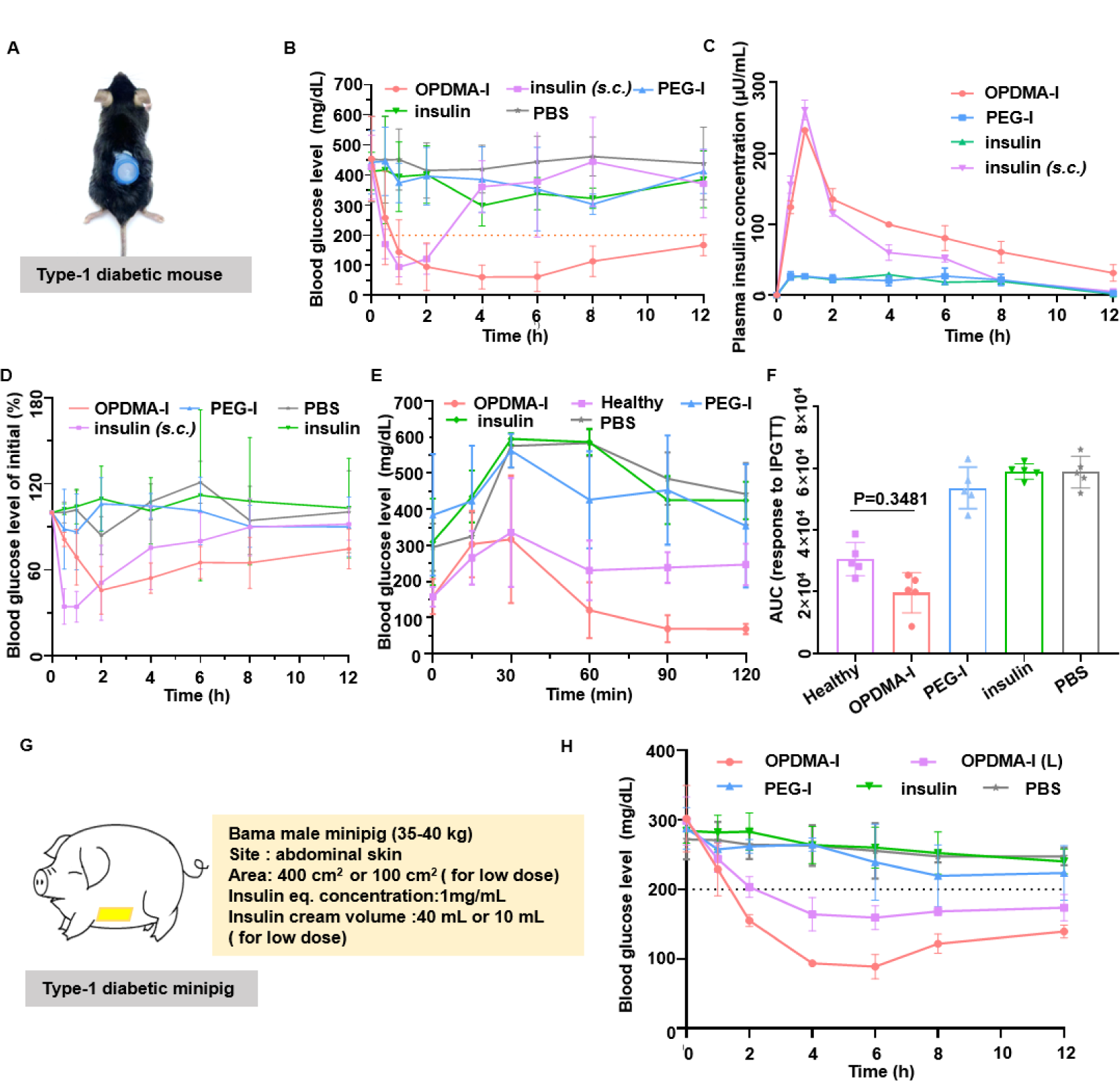
Hypoglycemic effect of OPDMA-I in STZ-induced diabetic mouse and minipig models. **A-C**, Transdermal treatment via a diffusion cell (1.77 cm^2^) on the dorsum skin of diabetic mice (**A)**, blood glucose levels **(B)** and blood insulin concentrations **(C)** of mice (n= 5) after transdermal treatment with PBS, native insulin, PEG-I, or OPDMA-I (0.2 mL, insulin-eq dose: 0.5 mg/mL, 116 U /kg); subcutaneous (*s.c*) injection of native insulin (5 U/kg) was used as a positive control. **D**, blood glucose levels of healthy mice after transdermal administration of OPDMA-I, PEG-I or native insulin, or *s.c* injection of insulin. **E**, *In vivo* intraperitoneal glucose tolerance test (IPGTT) in diabetic mice (n= 5) receiving transdermal treatments upon intraperitoneal administration of glucose at 1.5 g/kg; healthy mice were used as control. **F**, The area under the curve (AUC) from 0-120 min of IPGTT in diabetic mice calculated with the baseline set at the 0-min blood glucose reading. **G**, Schematic of an STZ-induced diabetic minipig treated with a topical cream containing OPDMA-I, PEG-I, or native insulin on the abdomen skin. **H,** blood glucose levels in diabetic minipigs (*n*= 3); insulin-eq concentration of 1mg/mL, 40 mL, application area: 400 cm^2^; for low dose group (L): 10 mL, application area: 100 cm^2^. Data are shown with mean ± SD.

Subsequently, we performed an intraperitoneal glucose tolerance test (IPGTT) at 1 h post-transdermal administration with a glucose dose of 1.5 g/kg to evaluate the glycemic regulation capacity of OPDMA-I. As shown in Fig. 1E, the BGLs of healthy mice gradually decreased within 1 h after a blood glucose peak of 330 mg/dL at 30 min post intraperitoneal glucose injection. The diabetic mice treated with OPDMA-I had a blood glucose profile similar to the healthy mice but with even lower BGLs after 1 h post-glucose injection. In contrast, the BGLs of the mice topically treated with insulin or PEG-I increased to even higher levels, ∼ 600 mg/dL, and maintained a hyperglycemic state over 120 min. The calculated area-under-the-curve from 0 to 120 min indicated that transdermal OPDMA-I had an excellent blood glucose regulation capability (Fig. 1F).

The porcine skin is considered a good human skin model in terms of its general structure, thickness, hair sparseness, and collagen and lipid composition^[17]^. Therefore, *in vivo* performance of OPDMA-I was further evaluated in an STZ-induced type-1 diabetic minipig model. OPDMA-I, PEG-I, or native insulin was dispersed in W/O cream and smeared on the abdomen skins of the diabetic minipigs. The blood glucose levels (BGLs) of the minipigs were monitored over time. (Fig. 1G). Similarly, the BGLs of the minipigs treated with the OPDMA-I cream decreased quickly to normoglycemia after 2 h with a minimum of 100 mg/dL at 6 h and maintained normoglycemic for over 12 h. In the case of a lower dose of OPDMA-I, the BGLs of the minipigs had a similar trend but with a slower decreasing rate and could also maintain normoglycemic levels with relatively higher BGLs. In contrast, the BGLs in minipigs treated with PEG-I or native insulin showed almost no significant reduction except for later due to fasting (Fig. 1H **and** Fig. S3B).

After 12 h-transdermal treatment with OPDMA-I, the stratum corneum (SC) of the treated skin sites in the mice and the minipigs maintained the micromorphological characteristics as the untreated skins, as observed by scanning electron microscopy (SEM) imaging (Fig. S4A). Therefore, OPDMA-I would not disrupt the SC barrier function. Furthermore, these skin sites were also not different from the controls, having negligible neutrophil infiltration and changes in the epidermal thickness and skin thickness (Fig. S4B) with barely any cell apoptosis (Fig. S4C). Similar results were confirmed in the porcine skin (Fig. S4D,E). There were also no observable adverse effects on the blood cell counts and biochemical measures, as well as on the liver and kidneys (Table S1).

### Skin permeability of OPDMA-I

The skin permeation of OPDMA-I was first measured and quantified using an *in vitro* three-dimensional skin equivalent EpiKutis^®^ model ^[22]^, and the corresponding permeation parameters were determined accordingly. The confocal laser scanning microscope (CLSM) imaging confirmed the quick permeation of OPDMA-I^FITC^ in the EpiKutis® (Fig. 2A). The steady-state flux (*J*_ss_) of OPDMA-I was significantly higher, and its permeability coefficient (*K*_p_) was more than nine-fold of that of insulin (Fig. 2B **and** Table S2).

**Fig. 2.**
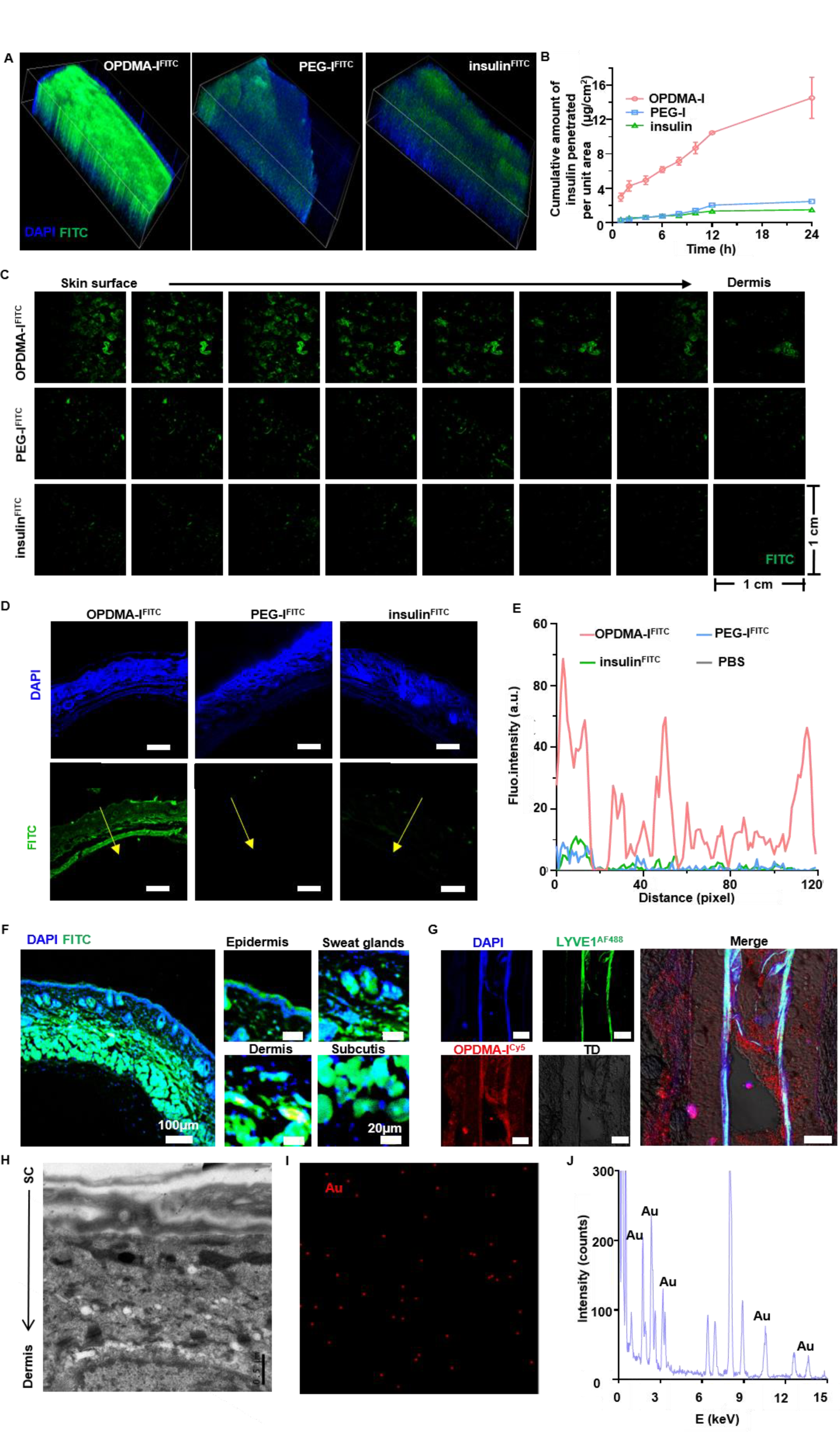
Skin permeability of OPDMA-I. **A,** 3D images of OPDMA-I^FITC^, PEG-I^FITC,^ or insulin^FITC^ distribution across the in vitro Three-Dimensional Skin Equivalent EpiKutis® model. (FITC-equivalent concentration, 1 μg/mL; volume, 0.2 mL; penetration area 0.081cm^2^; Width:1272.79 μm; Height: 1272.79 μm; Depth:170.00 μm). **B,** Permeation curves of OPDMA-I, PEG-I, and native insulin across EpiKutis® model. (insulin eq. concentration :0.5 mg/mL; volume:0.2 mL; penetration area: 0.081 cm^2^; n=3; data are shown as mean ± SD). **C,** Sequential images of OPDMA-I^FITC^ treated mice skin samples (FITC-eq. concentration, 1 μg/mL, application area: 1.77cm^2^). **D**, CLSM images of mice dorsal skin slices after 4 h penetration of 0.2 mL FITC-labeled OPDMA-I, PEG-I, and native insulin (FITC-eq. concentration, 1 μg/mL, application area: 1.77 cm2), 3 images obtained. Scale bars, 50 μm. **E,** FITC fluorescence intensity from the skin surface SC to the subcutis plotted along randomly selected lines (yellow arrow in **D**). **F**, The CLSM image of full-thickness mice dorsal skin at 4 h-post treatment of 0.2 mL OPDMA-I^FITC^. (FITC-eq.concentration:1 μg/mL, application area:1.77cm^2^).; Scale bar, 100μm. Stratum corneum (SC), the viable epidermis (VE), dermis, hair follicles, and subcutis within cryosection of excised full-thickness mice skin. **G,** Immunofluorescence staining of LYVE-1^AF488^ (green) in the subcutaneous tissue of rats at 4 h-post transdermal administration of 0.2 mL OPDMA-I^Cy5^ (red). (Cy5-eq.concentration:1 μg/mL, application area:1.77cm^2^) . Nuclei were labeled with DAPI (blue). Scale bars, 50 μm. **H,** Transmission electron microscopy (TEM) images of mice dorsal skin vertical to the skin’s surface 4 h-post treatment with 0.2 mL OPDMA-AuNPs (OPDMA-equivalent concentration: 0.5 mg/mL, application area: 1.77 cm^2^, n= 3, Scale bar, 0.5 μm.). **I,J,** Scanning transmission electron microscopy (STEM) and energy dispersive X-ray spectroscopy (EDS) analysis of the OPDMA-AuNPs treated skin cryosection. **I,** STEM elemental mapping images. **J,** EDS analysis element profiles.

Upon transdermal application, Cy5-labeled OPDMA-I (OPDMA-I^Cy5^) quickly adsorbed on the skin, displaying strong fluorescence intensity after 0.5 h post-treatment compared with the weak fluorescence of the mice skin treated with PEG-I^Cy5^ or insulin^Cy5^ (Fig. S5A). The permeation of FITC-labeled OPDMA-I (OPDMA-I^FITC^) in the inner layer of the mouse skin was further analyzed by sequential z-stack imaging using intravital two-photon microscopy. Figure 2C shows that after 4 h-transdermal application, OPDMA-I^FITC^ penetrated much deep into the skin tissues than PEG-I^FITC^ or insulin^FITC^, with more intense fluorescence at any depth. The transdermal OPDMA-I^FITC^could reach the tissue beneath the skin, suggesting the excellent skin penetration capability of OPDMA-I ^FITC^, while PEG-I^FITC^ or insulin^FITC^ could barely adsorb on the skin surface, let alone penetrate the skin tissues (Fig. 2D,E). *Ex vivo* fluorescence imaging indicated that the topically applied OPDMA-I^Cy5^ entered the circulation and accumulated mostly in the liver (Fig. S5B).

Confocal microscopy imaged the vertical section of the skin receiving 4 h-transdermal treatment with OPDMA-I^FITC^ (Fig. 2F) and its enlarged SC regions, the viable epidermis (VE), dermis, and hair follicle (Fig. 2G). OPDMA-I^Cy5^ distributed across the whole skin layers; most already reached the dermis layer, and some entered the lymphatic vessel labeled with LYVE1^AF488^. In order to further assess the skin penetration ability of OPDMA, we tethered OPDMA-SH on gold nanoparticles of 5 nm (OPDMA-AuNPs) and transdermally applied them to the mouse dorsum skin for 4 h. Transmission electron microscopy (TEM) (Fig. 2H), scanning transmission electron microscopy elemental mapping (Fig. 2I), and energy dispersive X-ray spectroscopy (EDS) (Fig. 2J) analysis showed that OPDMA-AuNPs appeared in the inter-corneocyte lipid lamella, viable epidermis, and dermis.

### Mechanism study of OPDMA-I penetrating the stratum corneum

The outer SC layers of mouse skin were peeled off for CLSM observation according to a previously reported method^[23]^ after 4 h-transdermal treatments with OPDMA-I^Cy5^, PEG-I^Cy5,^ or insulin^Cy5^. The SC layers from the OPDMA-I^Cy5^-treated skin had much stronger fluorescence than the other groups (Fig. S6A), and the fluorescence mainly surrounded the periphery of the corneocytes (Fig. 3A). TEM also revealed that OPDMA-AuNPs appeared at the inter-corneocyte lipid lamella (Fig. 3B). The OPDMA-I^FITC^ distribution in SC layers imaged by intravital two-photon microscopy confirmed the phenomenon (Fig. 3C **and** Fig. S6B). These results suggest that OPDMA-I crossed the skin SC through the para-corneocyte route. These results were also verified using quartz crystal microbalance to measure the adsorption of OPDMA-I on a model SC lipid membrane with the same composition (Fig. S7A).

**Fig. 3.**
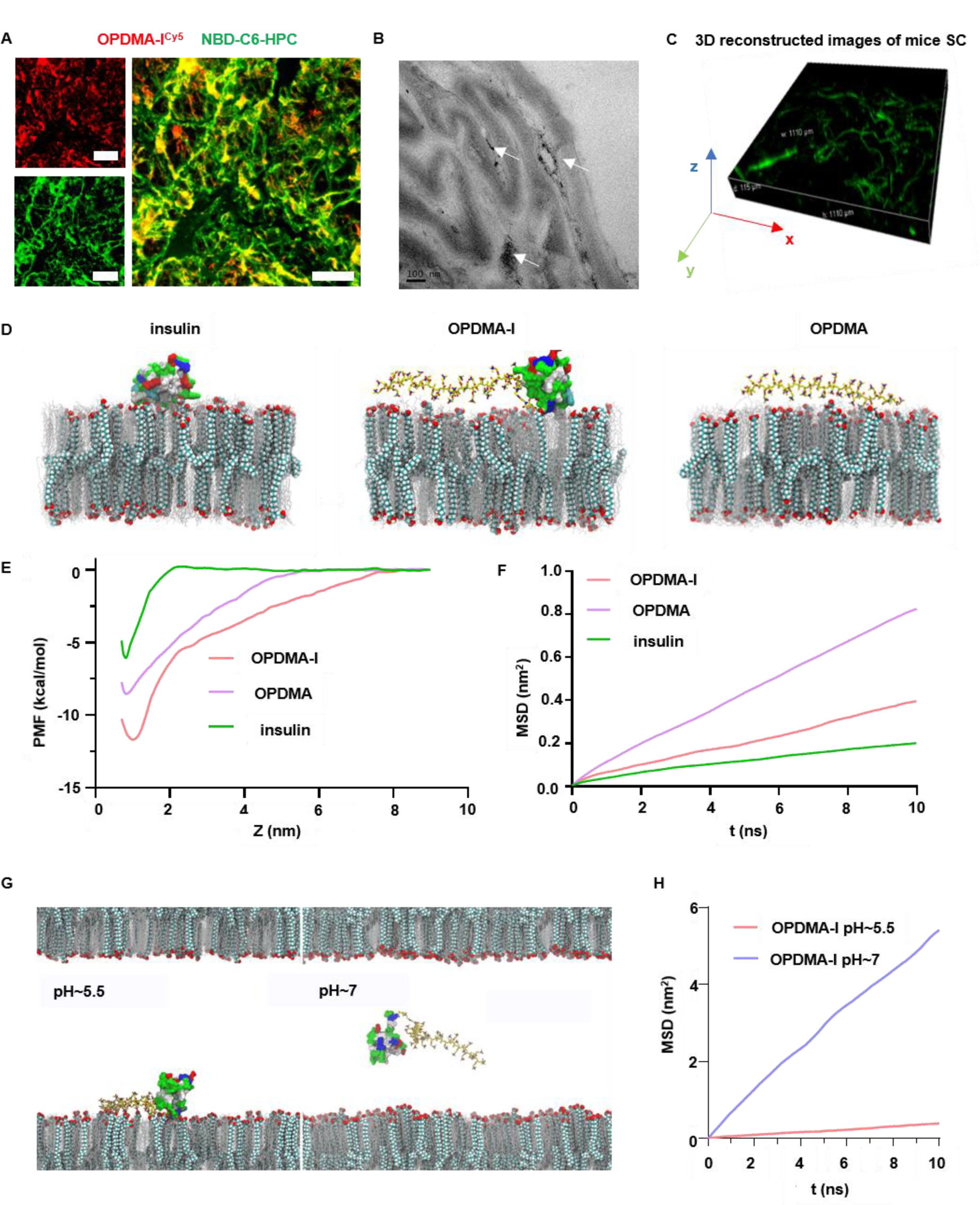
Mechanism study of OPDMA-I penetrating the stratum corneum. A, Fluorescence distribution of OPDMA-I^Cy5^ (red) in NBD-C6-HPC stained SC intercellular lipid (green) imaged by CLSM. SC samples were extracted from the deep layer of mouse dorsal skin by adhesive tape after in vivo skin permeation for 4 h-post transdermal application of 0.2 mL OPDMA-I^Cy5^. Cy5-eq. concentration, 1 μg/mL, application area: 1.77cm^2^. Scale bars, 25 μm. **B,** TEM images of mice skin vertical to skin surface after 4 h-treatment with OPDMA-AuNPs. OPDMA-eq. concentration: 0.5 mg/mL, 0.2 mL, application area: 1.77 cm^2^, n= 3. Scale bars, 100nm. **C,** The 3D reconstructed view of sequential images of OPDMA-I^FITC^ treated mice SC layers. **D,** Representative binding modes of insulin, OPDMA, or OPDMA-I on SC lipids. **E,** Potential mean force (PMF) results indicating the binding free energies of insulin, OPDMA, or OPDMA-I with SC lipids. Z represented the distance between the center of mass of insulin/OPDMA/OPDMA-I and the surface of the SC lipids. **F,** Mean squared displacement (MSD) results comparing the diffusivities of insulin, OPDMA, or OPDMA-I on SC lipids at pH 5.5. **G,** Representative interaction modes of OPDMA-I on the SC lipids at pH 5.5 and pH 7.0. **H,** MSD results comparing the diffusivities of OPDMA-I on SC lipids at the weak acid condition (pH 5.5) and at the neutral condition (pH 7.0).

All-atom molecular dynamics (MD) simulations were performed to investigate the interactions between insulin, OPDMA, or OPDMA-I with SC lipids in order to gain a deeper understanding of the underlying molecular mechanism behind OPDMA-I’s penetration (Fig. 3D-H). Their binding processes onto SC lipids at the weak acidic condition (pH 5.5) are displayed in **Movie S1**, showing that OPDMA and OPDMA-I adsorbed more rapidly onto SC lipids than insulin. The corresponding binding configurations are illustrated in Fig. 3D **and** Fig.S7B. Their binding free energies estimated by the potential of mean force (PMF) analyses (Fig. 3E) showed that the binding affinities of OPDMA (−8.5 kcal/mol) and OPDMA-I (−11.7 kcal/mol) with SC lipids were stronger than that of insulin (−6.0 kcal/mol). Interestingly, although OPDMA possessed a higher binding affinity with SC lipids than insulin, the decay of the binding free energy of OPDMA with the distance from the SC surface was much slower than that of insulin. This suggested that the SC membrane provided much lower “local trapping” (restriction in motion) for OPDMA than that of insulin, which should benefit faster diffusion of OPDMA on SC lipids. The MD simulations demonstrated the stronger diffusivity of OPDMA and OPDMA-I on SC lipids than insulin (Fig. S7C **and Movie S2**), which was also quantitatively verified by the mean squared displacements (MSD) calculations (Fig. 3F). PMF analyses along the surface of SC lipids (in the *X* direction) were performed to quantitatively estimate the energy barriers for the diffusions (Fig. S7D), which was 0.9 ± 0.1 kcal/mol for OPDMA, 2.2 ± 0.3 kcal/mol for insulin, and 1.7 ± 0.2 kcal/mol for OPDMA-I. Accordingly, these MD results suggested that OPDMA and OPDMA-I suffered less local trapping and diffused much faster than insulin on SC lipids. The MD simulations further compared the interactions of OPDMA-I with SC lipids at weak acidic (pH 5.5) and neutral conditions (pH 7.0) (Fig. 3G **and Movie S3)**. The electrostatic interaction between OPDMA and SC lipids became negligible at pH 7.0 as OPDMA became zwitterionic. Thus, OPDMA-I bound the SC lipids more weakly and diffused faster (Fig. 3H) at the neutral condition than that at the acid condition.

### The mechanism of OPDMA-I penetration of the viable epidermis

The permeability of OPDMA-I in the viable epidermis was first observed using 3D-cultured multilayer human immortal keratinocyte (HaCat) spheroids^[24]^. Despite being incubated with PEG-I^FITC^ or insulin^FITC^ for 12 h, the spheroids exhibited weak fluorescence both on their surface and interior (Fig. 4A). In contrast, after only 1 h of treatment with OPDMA-I^FITC^, the spheroids displayed a remarkably bright fluorescent surface and interior, which was further confirmed by linescan analysis (Fig. 4B). Surprisingly, the endocytosis and exocytosis inhibitors did not inhibit the OPDMA-I^FITC^ infiltration in the HaCat spheroids (Fig. S8A,B). Notably, OPDMA-I^Cy5^ remained exclusively localized on the cell membranes even after 24 h-incubation without any detectable cytoplasmic uptake (Fig. 4C **and** Fig. S8C). Indeed, the nanovesicles formed by the isolated cell membrane exhibited strong fluorescence, as shown in Fig. S8D,E. These findings suggest that OPDMA-I^FITC^ was unable to enter the cells and therefore did not infiltrate the HaCat spheroids via the transcellular route.

**Fig. 4.**
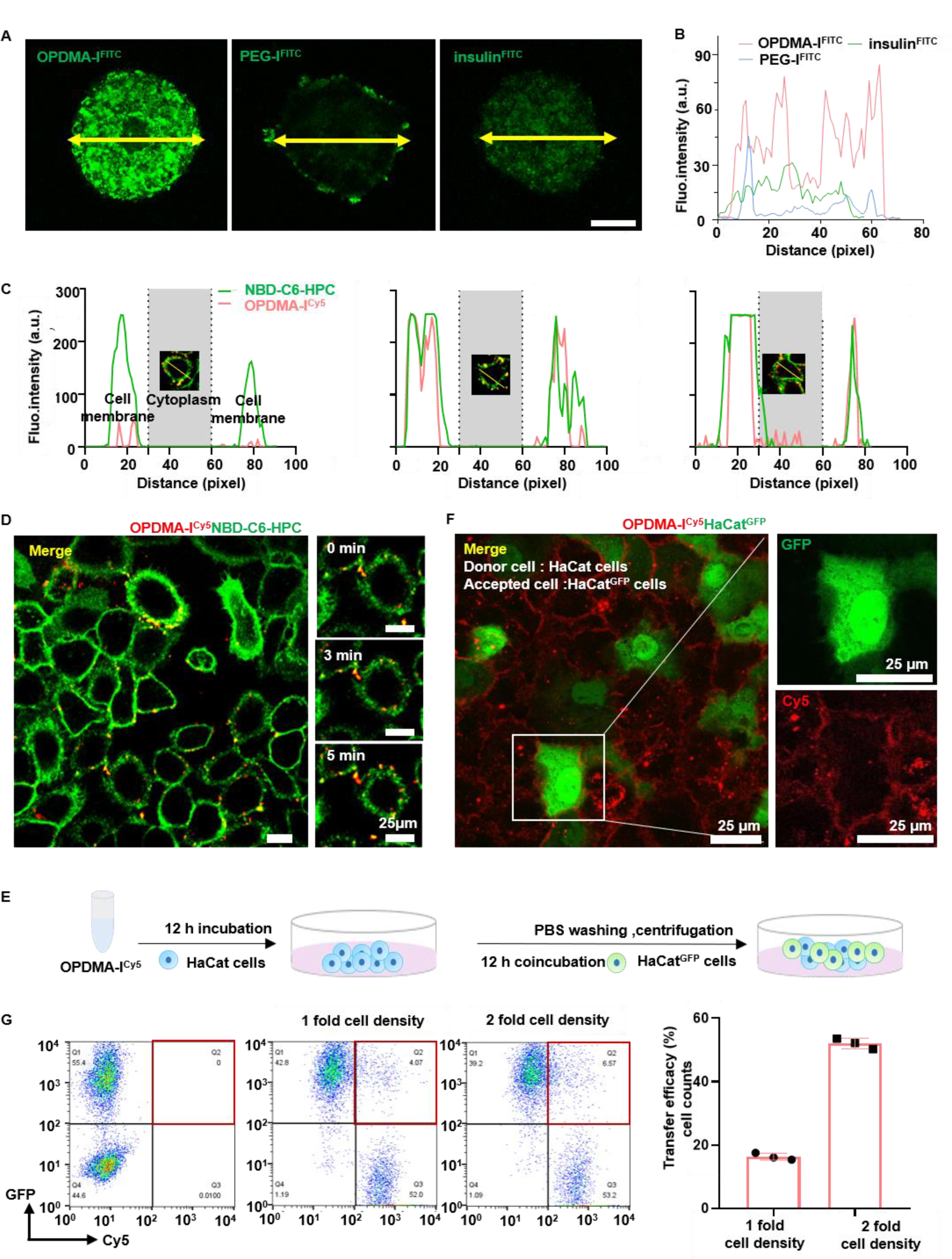
The mechanism of OPDMA-I penetration of the viable epidermis. **A-B,** 3D-cultured multilayer HaCat spheroids were separately pretreated OPDMA-I^FITC^, PEG-I^FITC,^ or insulin^FITC^ for 12 h and then imaged with CLSM by Z-stack tomoscan at 20 μm intervals. **A,** Representative images of FITC fluorescence distribution in the middle layers of spheroids. Scale bars, 100 μm. (3 images obtained) **.B,** FITC fluorescence intensity along the yellow arrows in **A. C,** Colocalization of NBD-C6-HPC-stained HaCat cell membrane (green line) and OPDMA-I^Cy5^ (red line) along the selected direction (yellow) in CLSM images of a randomly selected HaCat cell after 6 h,12 h, and 24 h incubation. **D,** CLSM images of OPDMA-I^Cy5^ jumping on the cell membranes. CLSM images of HaCat cell membrane adsorbing OPDMA-I^Cy5^ and fixed views obtained from Movie S4 in a time-lapse-acquisition mode; Cy5-eq. dose, 0.1 μg/mL. The cell membranes were stained with NBD-C6-HPC (green). Scale bars, 20 μm. **E,** Experiment illustration of direct transfer of OPDMA-I^Cy5^ ( Cy5 eq. dose: 0.1 μg/mL). **F,** Representative CLSM images of OPDMA-I^Cy5^ transfer from pretreated HaCat to untreated HaCat^GFP^ cells. Scale bar, 50μm. **G,** Flow cytometry analysis of the direct transfer of OPDMA-I^Cy5^ from HaCat to HaCat^GFP^.

CLSM imaging in a time-lapse-acquisition mode visualized the “jumping” of OPDMA-I^Cy5^ among the adjacent cells (Fig. 4D **and Movie S4)**. A cell-to-cell transfer assay was performed by mixing OPDMA-I^Cy5^-pretreated HaCat cells (donor cells) with HaCat cells stably expressing green fluorescence protein (GFP) (HaCat^GFP^, recipient cells) for co-incubation (Fig. 4E). CLSM imaging and flow cytometric analysis demonstrated that HaCat^GFP^ cells gradually obtained Cy5-fluorescence, and increasing the cell densities facilitated the OPDMA-I^Cy5^ transfer among cells (Fig. 4F,G). However, when the two cell populations were cultured separately on different coverslips but still maintained in the same medium, the transfer of OPDMA-I^Cy5^ was significantly inhibited (Fig. S8F,G). Therefore, OPDMA-I penetrated the viable epidermis by hopping on the membranes of adjacent cells rather than through transcytosis, which is observed in OPDMA and its micelles.^[25]^.

## Discussion

Noninvasive transdermal insulin delivery is undoubtedly an ideal alternative to hypodermic injection for diabetic management, yet it remains unsuccessful due to the unmet issues of safety, delivery efficiency, and reliability ^[26]^. Here we reported an OPDMA-conjugated insulin, OPDMA-I, capable of efficient, safe, and noninvasive transdermal insulin delivery, with glycemic regulation performance comparable to *s.c.* injected insulin. The synthesis of OPDMA-I was facile and efficient, similar to protein pegylation (Fig. S1). OPDMA-I maintained the same biological conformation and glycemic regulation capability as native insulin (Fig. S2D,E).

Transdermal treatment with OPDMA-I showed an efficient *in vivo* regulation of glucose levels in STZ-induced diabetic mice, even superior to *s.c.* injected insulin. Once applied to the mouse dorsal skin, OPDMA-I quickly penetrated the skin into the plasma, reaching the maximum insulin concentration at 1 h and maintaining considerable insulin levels for 12 h (Fig. 1C). The OPDMA-I that entered blood circulation mainly accumulated in the liver (Fig. S3D), the main organ where insulin exerts its glycemic regulation effects^[27]^. Furthermore, transdermal OPDMA-I lowered the BGLs as fast as the *s.c.* injected insulin and offered prolonged normoglycemic regulation of glucose levels, with no issues of hypoglycaemia (Fig. 1B). The IPGTT results indicate that transdermal OPDMA-I could efficiently defuse the glucose challenge, resulting in better blood glucose regulation in diabetic mice than healthy mice (Fig. 1E). Based on the encouraging findings in the diabetic mouse model, studies were further conducted on diabetic minipigs, showing that transdermal OPDMA-I was able to maintain minipig glucose levels in a normal range for over 12 h (Fig. 1H). Lowering the dose of OPDMA-I could still control blood glucose to normal levels within 2 h and keep the normoglycemia for 12 h. Moreover, noninvasive permeation of OPDMA-I through the skin holds great advantages over chemical or physical enhancing approaches and microneedles. OPDMA-I-treated mouse skin had no observable structural change in the SC microstructures, with no shedding of corneocytes and no widening of the intercellular gaps (Fig. S4A). Treatments with OPDMA-I also caused no noticeable inflammation or cell death in the skin of the mice and minipigs (Fig. S4B-E).

The transdermal process of OPDMA-I included three steps, (i) adsorption and enrichment on the acidic sebum, (ii) SC penetration *via* diffusion within the inter-corneocyte space, and (iii) diffusion from the viable epidermis to the dermis *via* hopping on the cell membranes.

Generally, molecules would penetrate the SC *via* three approaches: the appendageal path, the para-corneocyte path, and the transcellular path (through corneocytes) ^[28]^. After transdermal application, OPDMA-I was mainly distributed in the inter-corneocyte lipids rather than inside the corneocytes (Fig. 3A-C), indicating that OPDMA-I possibly penetrated the SC through para-corneocyte gaps.

MD simulations confirmed the stronger affinity towards the SC lipids of OPDMA-I and enhanced diffusivity in SC compared with insulin. The enhanced binding affinity was primarily due to the strong electrostatic interaction between the positively charged OPDMA and negatively charged fatty acid at the acidic pH. This electrostatic interaction was long-ranged and decayed much slower with the increasing distance from the SC surface than the short-ranged van der Waals (vdW) interaction that dominates the interaction between insulin and SC lipids (insulin carries the same net negative charge with SC lipids at pH 5.5) (Fig.3E). Therefore, the interaction of OPDMA with SC lipids was less sensitive to the surface roughness of lipid membrane than that of insulin (Fig. S7B), thus encountering lower energy barrier during diffusion (Fig. S7D). More importantly, as thermal fluctuation is the driving force for diffusion, the strong fluctuation of OPDMA provides OPDMA-I an additional driving force to overcome its local energy barrier, thus leading to faster diffusion of OPDMA-I on the SC lipids than insulin **(Movie. S2)**. Furthermore, the increasing pH in the deep SC layers reduced the protonation of OPDMA, weakening its binding and promoting its diffusion (Fig. 3H **and Movie S3)**.

The subsequent diffusion of OPDMA-I was in the viable epidermis and dermis. Our previous work demonstrates that OPDMA and its micelles efficiently permeated tumor spheroids via transcytosis^[25]^. However, in this case, OPDMA-I^Cy5^ could adhere to the HaCat cell membrane but not enter the cytoplasm even after 24 h incubation (Fig. 4C **and** S8C). OPDMA-I^FITC^ could also infiltrate the 3D HaCat cell spheroids mimicking viable epidermis, but its permeability was not inhibited by endocytosis and exocytosis inhibitors (Fig. S8A,B). Therefore, a paracellular rather than transcellular pathway (transcytosis) was involved in OPDMA-I’s penetration in the viable epidermis. OPDMA-I^Cy5^ could be transferred among adjacent cells; Increasing the cell density promoted the delivery efficiency (Fig. 4E-G), while separating the cells in a co-culture medium significantly inhibited this intercellular transfer of OPDMA-I^Cy5^ (Fig. S8F,G). Further studies revealed that OPDMA-I hopped from one cell membrane to another **(Movie S4)**, indicating that OPDMA-I penetrated the viable epidermal via hopping among adjacent cells.

In summary, this work demonstrated the first completely noninvasive transdermal delivery of insulin that exerts an *in vivo* hypoglycemic effect for diabetic treatment as effectively as *s.c.* insulin injection. The reversible binding of OPDMA with SC lipids along the characteristic pH gradient of the skin accounted for the fast skin penetration of OPDMA-I. The acidity of sebum protonated OPDMA, facilitating its binding and enrichment on the SC. Conversely, increased pH in deeper layers of the SC liberated OPDMA-I from lipids, allowing for diffusion into the epidermis. OPDMA-I did not enter viable epidermis and dermis cells but hopped on the cell membrane, which avoided intracellular degradation of OPDMA-I and facilitated rapid penetration. Together, the transdermal application of OPDMA-I resulted in an immediate hypoglycemic effect. Importantly, this OPDMA-modification strategy may not be limited to transdermal insulin delivery but can also be used to deliver other biomacromolecules, including bioactive peptides, proteins, and nucleic acids.

## Acknowledgments

This work was supported by the National Key Research and Development Program (2021YFA1201200) and the National Natural Science Foundation (51833008) of China, and the Zhejiang Key Research Program (2020C01123).

## Materials and Methods

### Cell lines and mice

HaCat cell lines were obtained from the Cell Bank of the Chinese Academy of Sciences. The cells expressing green fluorescence protein (HaCat-GFP cells) were established by lentivirus transfection of the GFP plasmids into HaCat cells according to the manufacturer’s protocol (Shanghai Genechem). All the cell lines were incubated in a nutritious DMEM medium (containing 10% FBS and 1% (v/v) penicillin-streptomycin) at 37 °C with 5% CO _2_.

C57 BL/6J male mice (6-8 weeks, 25 g) were purchased from Shanghai Slaccas Laboratory Animal. The mice were housed in the Zhejiang Academy of Medical Sciences under specific pathogen-reduced conditions.

Guangxi Bama-minipigs (male, 6 months old, 35-40 kg) were purchased from Shanghai Jiagan Laboratory Animal and housed in the Laboratory Animal Center of Zhejiang University.

### Animal use statement

All animal experiments were carried out according to the protocols approved by the Zhejiang Academy of Medical Sciences and the First Affiliated Hospital of Zhejiang University.

### Experimental materials

If not otherwise indicated, all materials were purchased from Sinopharm Chemical Reagent Co., Ltd. (Shanghai, China). Dichloromethane (CH_2_Cl_2_, DCM) and tetrahydrofuran (THF) were distilled over calcium hydride (CaH_2_) or treated with a 4 Å molecular sieve. Trifluoroacetic acid (TFA), 2,2′-bipyridine, 1-dodecanethiol (98%), 2-bromo-2-methylpropionic acid (98%), carbon disulfide (anhydrous, 99%), potassium phosphate (98%), azobisisobutyronitrile (AIBN) *N*-hydroxysuccinimide (NHS), *N,N’-*dicyclohexylcarbodiimide (DCC) and 4-*N,N-* dimethylaminopyridine (DMAP) were purchased from Energy Chemical (Shanghai, China). Fluorescein isothiocyanate (FITC) and cyanine-5 succinimidyl ester (Cy5-NHS) were purchased from Lumiprobe Corporation (Shanghai, China). Human recombinant insulin (29 U/mg) was purchased from Solarbio LIFE SCIENCES (Beijing, China). NBD-C6-HPC was purchased from J&K Scientific (Beijing, China). 2-(Dodecyltrithiocarbonate)-2-methylpropionic acid (DDMAT) and 6-(maleimido)hexanoic acid succinimidyl ester (BMPS) were purchased from Aladdin (Shanghai, China). 2-Iminothiolane hydrochloride (Traut’s reagent) and streptozotocin (STZ) were purchased from Macklin (Shanghai, China). Gold nanoparticles (AuNPs, 5nm) were purchased from XFNANO (Nanjing, China)

### Synthesis of N-[2-(N-tert-butoxycarbonylamino)]ethyl-2-(dodecyltrithio-carbonate)-2-methylpropionamide (TAC)

DDMAT (2 mmol), DCC (2.5 mmol), DMAP (3 mmol), and NHS (2 mmol) were dissolved in 20 mL CH_2_Cl_2_ and stirred at room temperature for 24 h. The mixture was filtrated and washed with 100 mL water, dried over MgSO_4_, filtered, and concentrated under reduced pressure to obtain the crude product. The crude product obtained above was dissolved in 20 mL CH_2_Cl_2_. TFA (2.5 mmol) and *N*-Boc-EDA (2.0 mmol) were added dropwise to the solution and stirred for 12 h. The mixture was successively washed twice with water and saturated brine, dried over MgSO_4,_ and filtered. The filtrate was concentrated under reduced pressure to obtain a crude product.

The crude product was passed through a column packed with silica gel using a mobile phase of *n*-hexane and ethyl acetate mixture (10:1). The yellow solid *N*-[2-(*N*-tert-butoxycarbonylamino)]ethyl-2-(dodecyltrithiocarbonate)-2 methylpropionamide (TAC) was obtained.

### Synthesis of Boc-amino-terminated poly[2-(N,N-dimethylamino)ethyl methacrylate] (N-Boc-PDMA)

DMA (30 mmol), TAC (0.5 mmol), and AIBN (0.05 mmol) were dissolved in THF (30 mL) in a Schlenk flask and bubbled with dry N_2_ for 30 min. The reaction was carried out at 65 °C for 12 h. After terminating the polymerization by opening the flask, the solution was concentrated and poured into cold n-hexane. The precipitated *N*-Boc-PDMA was isolated and then dried under a vacuum.

### Synthesis of Boc-amino-terminated poly[2-(N-oxide-N,N-dimethylamino)ethyl methacrylate) (N-Boc-OPDMA)

*N*-Boc-PDMA (0.5 g) was dissolved in 5 mL of 30% hydrogen peroxide (H_2_O_2_) solution. The mixture was stirred at room temperature for 4 h and then dialyzed against deionized water to remove the unreacted H_2_O_2_ completely. *N*-Boc-OPDMA was obtained after lyophilization.

### Synthesis of OPDMA-NH_2_

*N*-Boc-OPDMA (500 mg) was dissolved in 5 mL CH_2_Cl_2,_ and TFA (5 mL) was added dropwise under ice-cooling. This solution was stirred for 2 h at room temperature. OPDMA-NH_2_ was obtained by precipitating with diethyl ether.

### Synthesis of OPDMA-BMPS

OPDMA-NH_2_ (2 mg) was dissolved in PBS (pH 7.4), followed by 6-(maleimido) hexanoic acid succinimidyl ester (BMPS, 2 eq). This solution was stirred for 2 h at room temperature. OPDMA-BMPS was obtained after repeated ultrafiltration (MWCO: 3.5 kDa).

### Synthesis of OPDMA-I

The free amine group of insulin was converted to sulfhydryl by 2-iminothiolane hydrochloride (Traut’s reagent). Traut’s reagent (0.0472 mg, 0.3 μmol) was added to the insulin solution (2 mg, 0.3 μmol, pH 7.4). This solution was stirred for 2 h at room temperature, and insulin with an SH group (insulin-SH) was obtained by ultrafiltration (MWCO: 10 kDa) and lyophilization.

The PBS buffer (pH 7.4) solutions of insulin-SH (2 mg, 0.3 μmol) and OPDMA-BMPS (2 mg, 0.3 μmol) were combined and stirred at room temperature for 2 h. OPDMA-I was obtained by ultrafiltration (MWCO: 10 kDa) and lyophilization.

### Synthesis of OPDMA-AuNPs

OPDMA-SH was prepared similarly to insulin-SH described above, except insulin was replaced by the equimolar OPDMA-NH_2_. OPDMA-SH (1 mg) was added to 1 mL AuNPs solution (carboxyl modification amount of 200-300 μmol/L) and stirred at room temperature for 2 h. The OPDMA-AuNPs were purified by ultrafiltration (MWCO: 10 kDa) 5 times.

### Labelling OPDMA-I, PEG-I, and native insulin with FITC or Cy5

FITC or Cy5-NHS solution in DMSO (5 mg/mL) was added dropwise with gentle stirring to insulin or its conjugate (10 mg/mL) in PBS (pH 7.4) with the molar ratio of FITC (or Cy5): insulin of 1:1 at room temperature under the dark overnight. The labeled insulin or its conjugate was purified by repeated ultrafiltration (MWCO: 3.5 kDa) until no free FITC or Cy5 fluorescence was detected in the filtrate.

### RP-HPLC chromatography

The HPLC was carried out on a 1525 binary HPLC pump with an ACE 5C18-300 column. Mobile phase A was 0.2 M sodium sulfate solution at pH 2.3 at 25 °C. Mobile phase B was acetonitrile at pH 2.3 at 25 °C. The injection volume was 20 μL, and the detection temperature was 30℃. The detection wavelength was 214 nm.

### CD spectroscopy

Far UV-CD spectra were recorded at 37 °C on a JASCO J-815 spectropolarimeter. Quartz cuvettes with a path length of 1 mm were used. Each spectrum was an average of four scans recorded from 250 to 190 nm at 1 nm steps.

### MALDI-TOF MS analysis

Prepared samples (2 μL), OPDMA-I, PEG-I, or native insulin solution, were spotted on the MALDI target plate equipped with the MALDI (Bruker Daltonics Inc.) system and then dried under an N_2_ stream. After the sample was dried, 1 μL of the matrix solution (DHB) was coated on each sample well. Mass spectrometry was performed in linear mode.

### In vivo studies using STZ-induced diabetic mice

The type-1 diabetic mice were induced by STZ infusion. Briefly, healthy male C57BL/6J mice (6-8 weeks; 25 g) were *i*.*p*. injected with streptozotocin (STZ, 150 mg/kg, dissolved in disodium citrate buffer (pH 4.5) at a concentration of 10 mg/mL). After 7 d of recovery, the successful establishment of the STZ-induced diabetic mice was confirmed by monitoring the glucose levels.

The diffusion cell was loaded with insulin or its conjugates and attached to the dorsal skin of the mice. The diffusing area was 1.77 cm^2^. The equivalent insulin dose for each mouse was 116 IU/kg (insulin equivalent concentration: 0.5 mg/mL, 0.2 mL per mouse). The blood glucose levels were measured from tail vein blood samples (∼3 μL) of mice using a calibrated Sinocare glucose meter.

Measurement of the plasma insulin concentrations: blood samples (25 μL) were drawn from the tail vein of mice at timed intervals. The plasma was isolated and stored at −20 °C. The plasma insulin concentration was determined using a Human Insulin ELISA kit according to the manufacturer’s protocol (Invitrogen). The relative bioavailability (RA) of transdermal administrated OPDMA-I, PEG-I, and insulin (I) was calculated.

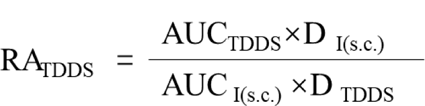

An Intraperitoneal glucose tolerance test was conducted to confirm the *in vivo* glucose regulation ability at 1 h after the administration of OPDMA-I. Briefly, mice were applied with the insulin products on the skin as described above. After 1 h, a glucose solution in PBS was intraperitoneally injected into the mice at a dose of 1.5 g/kg. The glucose levels were monitored over time. The glucose tolerance test on healthy mice was used as a control.

### OPDMA-I^Cy5^ skin adsorption and biodistribution in male C57BL/6J mice

Male C57BL/6J mice were randomly grouped (three mice per group). Each mouse was applied with 0.2 mL OPDMA-I^Cy5^, PEG-I^Cy5,^ or insulin^Cy5^ solution (Cy5-eq.dose, 1μg/mL) on the mouse dorsal skin with an area of 1.77 cm^2^. The mice were sacrificed at 0.5 h, 1 h, 2 h, 4 h, or 8 h after administration. The fluorescence intensities in the mouse dorsal skins were measured by an *in vivo* imaging system (IVIS Lumina XRMS Series III, PerkinElmer). The organs and tissues, including the heart, liver, spleen, lung, and kidney, were also harvested, and their fluorescence intensities were measured using the imaging system.

### In vivo studies using STZ-induced diabetic minipigs

Three male Guangxi Bama Mini-Pigs aged 6 months (weight: ∼35-40 kg) were used. Diabetes was induced in the minipigs by STZ infusion. STZ (75 mg/mL) in freshly prepared disodium citrate buffer (pH 4.5) was infused into the minipigs (150 mg/kg) within 10 min. The glucose levels were monitored using the CGMS^[17]^ (Dexcom G4 Platinum Continuous Glucose Monitor, Dexcom). After 7 days of recovery, the blood glucose levels constantly higher than 250 mg/dL indicate the successful establishment of the insulin-deficient diabetes model.

OPDMA-I, PEG-I, or native insulin was dispersed in cream (W/O). The oil phase was composed of stearic acid, lanolin, and vaseline; the water phase was composed of glycerin, triethanolamine, and ethylparaben). The insulin-equivalent concentration was 1 mg/mL. The cream was topically applied on the minipigs’ abdominal skin at an insulin-equivalent dose of 40 mg (cream volume: 40 mL, application area: 400 cm^2^) or 10 mg (cream volume: 10 mL, application area: 100 cm^2^). The cream was fixed on the skin using plastic wrap. The blood glucose levels of minipigs were continuously monitored using CGMS.

### In vitro and in vivo biocompatibility tests

Histological Assay: The skin of diabetic mice and minipigs after the treatments were fixed in 4% paraformaldehyde and embedded in paraffin. The skin samples were sectioned into 5 μm-thick slices, each subjected to hematoxylin and eosin (H&E) staining and examined under an upright microscope. The apoptotic cells in the skin were identified using a terminal deoxynucleotidyl transferase (TdT)-mediated dUTP nick end labeling (TUNEL) apoptosis assay kit and analyzed using CLSM.

Routine blood test and blood chemistry analysis: The *in vivo* serum and organ toxicology of transdermal OPDMA-I in minipigs were evaluated via the standard blood test and blood chemistry analysis. Blood samples were analyzed at the School Hospital of Zhejiang University.

### In vivo skin permeation

Four hours after topical administration of OPDMA-I^FITC^, PEG-I^FITC,^ or insulin^FITC^ (FITC-eq. concentration, 1μg/mL; 0.2 mL per mouse), the mice were euthanized, and their full dorsal skins were obtained. The skins were carefully washed with PBS three times and fixed with 4% PFA. The skin samples were sectioned into 10 μm-thick slices using a cryostat (UV800, Leica Microsystems, Bannockburn, IL). The slices were stained with DAPI for labeling cell nuclei.

The skin slices were imaged by CLSM with excitation at 405 nm for DAPI and 488 nm for FITC. The CLSM images were analyzed using ImageJ software.

### Skin penetration analysis by IntraVital Two-photon Microscopy

The mice were topically treated with OPDMA-I^Cy5^, as mentioned above. The skin was mounted between coverslips and slide glasses for two-photon imaging analysis. The laser wavelength for two-photon excitation was 480 nm, and the laser power delivered to the skin sample was 90 mW. Sequential z-stack images were captured at 3 μm intervals from the skin surface until the fluorescence signal became undetectable. The XZ-axis orthogonal view of the stratum corneum layers was reconstructed using ’volume viewer’ plugins in ImageJ.

### TEM imaging and STEM-EDS analysis of OPDMA-AuNPs-treated skin cryosections

C57BL/6J male mice were topically administrated with 0.2 mL OPDMA-AuNPs (OPDMA-eq. concentration 0.5 mg/mL). After 4 h, the mice were anesthetized and perfused with fixative (containing 4% PFA and 2.5% glutaraldehyde) by cardiac injection to fix the capillaries in the dermis. The skin was collected and fixed overnight at 4 °C. The samples were trimmed into small sections and then treated with 1% osmium tetroxide (diluted in 100 mM cacodylate buffer) for 2 h, dehydrated with ethanol and acetone, and embedded in Spurr resin. Ten-μm-thick slices were sectioned (LKB 11800 Pyramitome) and stained with toluidine blue to select the desired regions. The ultrathin sections (50-100 nm) were prepared using an ultramicrotome (UC7, Leica) and then stained with lead citrate and uranyl acetate for observation (H-7650, Hitachi).

STEM-EDS analysis of the skin cryosections was performed on a Titan Chemi STEM electron microscope at an accelerating voltage of 200 kV equipped with an EDS detector.

### Stratum corneum sample collection and observation

The procedure followed the method reported in the literature.^[23]^ Four hours after topical administration of OPDMA-I^Cy5^, PEG-I^Cy5,^ or insulin^Cy5^ (Cy5-eq. concentration 1 μg/mL, 0.2 mL per mouse), the treated sites were washed three times with PBS and dried carefully. A piece of double-sided adhesive tape (Scotch 3M, St Paul, MN, USA) was pressed on the skin surface for 2 seconds and then peeled off in the longitudinal direction. The middle part of the tape was fixed on a microscope slide and stained by NBD-C6-HPC, then observed under CLSM at 488 nm excitation for NBD-C6-HPC and 647 nm excitation for Cy5.

### Quartz crystal microbalance with dissipation (QCM-D) experiments

Model SC lipid membranes were deposited and annealed on quartz crystals using the reported procedure ^[29]^. Briefly, the above solution was dropped to the sensor surface and settled naturally until solvent evaporation to form a thin layer of model SC membrane on the sensor, dried under vacuum overnight, and annealed at 70 °C for 10 min.

The QCM-D measurements were performed with a Q-Sense E4 unit (Q-Sense AB, Sweden). The quartz sensor coated with the SC lipid layer was carefully mounted in the flow module with the active surface facing the testing solutions or suspensions. The flow module was mounted on the chamber platform, and the solution was pumped into the flow module with an IPC-N peristaltic pump (Ismatec, Switzerland) at a flow rate of 150 Μl/min. All QCM-D experiments were carried out at 25.00 ± 0.02℃. The frequency and dissipation shifts were continuously recorded during the experiment.

### OPDMA-I transportation across three-dimensional skin equivalent EpiKutis®

A three-dimension skin model (EpiKutis®) was fabricated according to the literature reported.^[30]^ Briefly, keratinocytes (5 x 10^5^) were seeded on the permeable membranes of transwell chambers, which were cultured at 37°C in a 5% CO _2_ atmosphere for 2 days, then cultured at the air-liquid interface for 8 days with daily medium replacement. Then a complete EpiKutis 3D model was obtained and used as a skin model for investigating OPDMA-I skin permeability. OPDMA-I, PEG-I, or insulin solution (insulin-eq. concentration: 0.5 mg/mL, 0.2mL) was added to the donor compartment, and 0.4 mL fresh medium was added to the receiving compartment. The temperature was maintained at 37 °C . At timed intervals, 50 μL solution was withdrawn from the receiving compartment, and an equal volume of fresh medium was added. The insulin concentration was quantified by an ELISA kit. The unit conversion was calculated according to the following formula, and the cumulative amount of insulin permeating per unit area of the model skin (Q_n_) was calculated:

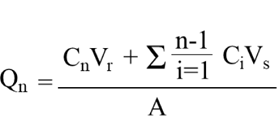

Where Q_n_ is the cumulative amount of insulin permeating per unit area (μg/cm^2^), C_n_ is the insulin concentration in the receiver cell at sampling time point t (μg/mL), C_i_ is the insulin concentration of the receiving liquid at the intermediate point (μg/mL), V_r_ is the volume of the receiving pool (0.4 mL), V_s_ is the volume of the sampled solution (50 μL), A is the effective transmission area of the EpiKutis® (0.081 cm^2^).

The steady-state flow rate (*J*_ss_) was obtained as the slope of the curve of Q_n_ as a function of time. The apparent permeability coefficient (*K*_p_) is calculated by the following formula:

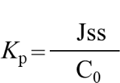

Where C_0_ is the initial insulin or its conjugate concentration in the donor cell.

EpiKutis® (0.081 cm^2^) was incubated with 0.2 mL OPDMA-I^FITC^, PEG-I^FITC,^ or insulin^FITC^ (FITC equivalent concentration, 1 μg/mL) for 4 h and then washed with PBS three times, stained with DAPI, and finally imaged using CLSM with the *Z*-stack tomoscan model at 10 μm intervals from the bottom to top of the 3D skin model with excitation at 405 nm for DAPI, and 488 nm for FITC.

### Molecular dynamics simulations

The insulin structure was taken from the Protein Data Bank (1AI0^[31]^, Chain I). The structures of OPDMA and OPDMA-I were constructed by Avogadro software^[32]^. The OPDMA was set to have 32 repeating units. The force field parameters of OPDMA and OPDMA-I were obtained from the Paramchem webser ver^[33]^ and CGenFF^[34]^. The SC lipid membrane, composed of an equimolar mixture of ceramide, cholesterol, and free fatty acids, was generated by the membrane builder of CHARMM-GUI^[35, 36]^. The CHARMM36^[37]^ force field was used to model the insulin and SC lipids. At pH 5.5, 20% of the *N*-oxide groups of OPDMA were protonated, whereas 50% of the fatty acids were deprotonated. At pH 7.0, OPDMA was zwitterionic, while all fatty acids were deprotonated. Each system was solvated in an 11.4 × 11.4 × 16.1 nm^3^ water box with ∼219000 atoms. Water molecules were modeled by the TIP3P water model^[38]^. Na^+^ and Cl^−^ ions were added to neutralize each system and bring its total ionic strength to the physiological concentration of 150 mM.

All MD simulations were carried out using the program GROMACS 2020.6^[39, 40]^. VMD^[41]^ was used for trajectory visualization. The covalent bonds with hydrogen atoms were constrained by the LINCS algorithm^[42]^, which allows a time step of 2 fs. The long-range electrostatic interactions were calculated using the particle-mesh Ewald method^[43]^, whereas the vdW interactions were calculated with a smooth cutoff of 1.2 nm. Periodic boundary conditions were applied in all directions. The NPT ensemble with semi-isotropic pressure coupling was applied with the pressure (1 bar) controlled by the Parrinello-Rahman barostat^[44]^ and the temperature (310 K) by the v-rescale thermostat^[45]^. Periodic boundary conditions were applied in all directions. Before production runs, the lipid system was equilibrated for 100 ns. Then 200-ns runs were conducted to monitor the adsorptions of insulin, OPDMA, and OPDMA-I on SC lipids. After adsorptions, three independent 100-ns runs were further performed for each of insulin, OPDMA, and OPDMA-I to monitor their diffusions on SC lipids. In production simulations, all atoms were free to move.

### Potential of mean force (PMF) analyses

The PMF results were calculated using the umbrella sampling protocol^[46]^. To estimate the energy barriers for diffusions of insulin, OPDMA, or OPDMA-I on SC lipids, the PMF setups were similar to the aforementioned MD setups. The total transverse distance along each representative path was 2 nm, which was divided into 20 windows with a resolution of 0.1 nm. Position restraints were applied to the SC lipids when the energy barriers were scanned. For monitoring the adsorptions of insulin, OPDMA, or OPDMA-I on SC lipids, the simulation boxes were extended to 11.4 × 11.4 × 22.1 nm ^3^ by adding 0.15 mM NaCl solution in the boxes. After a further 30-ns equilibration, the sampling path of each system was obtained by pulling insulin, OPDMA, or OPDMA-I in the perpendicular (*Z*) direction to the SC lipid membrane with a constant velocity of 0.2 nm/ns. The total sampling distance in the *Z* direction was 9 nm for each system, which was divided into 90 windows with a resolution of 0.1 nm. At each window, the system was first equilibrated for 5 ns, followed by a 35-ns productive umbrella sampling with a restraint force constant of 1000 kJ /mol/ nm^2^.

### Viable epidermis penetration study using 3D-cultured multilayer HaCat spheroids

Multilayer cell spheroids were prepared using the hanging-drop method. HaCat cells were suspended in fresh DMEM medium (containing 0.12% w/v methylcellulose) at a density of 4×10 ^5^ cells per milliliter. The cell suspensions (25 μL) were dropped on the lids of the cell culture plate to form uniform droplets, and 20 mL PBS was added to the plate to keep the droplets moist. The cells were incubated for 72 h and formed dense spheroids, which were transferred to an agarose-coated (1% w/v in PBS) 96-well plate with one spheroid per well and incubated for another 72 h to mature. The spheroids were incubated with OPDMA-I^FITC^, PEG-I^FITC,^ or insulin^FITC^ at a FITC-equivalent dose of 0.1 μg/mL for timed intervals. The spheroids were washed with PBS and imaged using CLSM by *Z*-stack tomoscan at 20 μm intervals from the bottom to the middle of the spheroids. The integration of FITC fluorescence density and linescan analysis was performed using ImageJ software.

For the effects of the endocytosis and exocytosis inhibitors on the penetration of OPDMA-I^FITC^, the HaCat spheroids were separately treated with PBS, wortmannin (2 μM), cytochalasin D (20 μM), monensin (20 μM), nocodazole (10 μM) or brefeldin A (10 μM) for 2 h, and then incubated with OPDMA-I^FITC^ (FITC-equivalent dose, 0.1 μg /mL) for 4 h. The HaCat spheroids were imaged and analyzed as described above.

### Transcellular delivery of OPDMA-I^Cy5^

HaCat cells were seeded on coverslips (i and ii) and incubated overnight. The cells on a coverslip (i) were first cultured with OPDMA-I^Cy5^ (Cy5-eq. concentration at 0.1μg/mL) for 4 h, then rinsed with PBS three times and co-incubated with fresh cells on the coverslip (ii) in fresh medium for 12 h. Afterward, the cells were washed with PBS, and the cell membrane was stained with NBD-C6-HPC (1 μM) for 5 min before imaging by CLSM at an excitation of 488 nm for NBD-C6-HPC and 640 nm for Cy5.

### Preparation and characterization of HaCat-derived nanovesicles

HaCat-derived nanovesicles were harvested by a reported method.^[47]^ Briefly, OPDMA-I^FITC^, PEG-I^FITC^, or insulin^FITC^ (FITC-eq. dose of 0.1 μg/mL)-treated HaCat cells were detached by scraping and resuspended in PBS (10^6^ cells/mL). The cell suspension was incubated in a 43 °C water bath for 1 h and recovered in a 37 °C incubator for 24 h. Subsequently, the cell suspension was sequentially extruded 11 times through 10, 5, 1, and 0.2 μm polycarbonate membrane filters via a mini extruder (Avanti Polar Lipids). The suspension was centrifuged at 1500 g for 30 min. Finally, the supernatant was centrifuged at 14 000 g for 1 h to collect HaCat-derived nanovesicles. Nanoparticle tracking analysis (NTA) was applied to measure particle size and concentration.

Purified nanovesicles were diluted in PBS to a final volume of 1 mL. The solution was loaded into the sample chamber of a NanoSight LM10 instrument (Malvern Panalytical, Malvern, UK). Five videos of 60 seconds each were recorded for each sample at 25 °C and a syringe speed of 40 µL/s. After capture, videos were analyzed with the in-build NanoSight Software NTA 3.1 Build 3.1.46. The fluorescence intensity of these nanovesicles was detected with a microplate reader excitation at 488 nm for FITC.

### Colocalization of OPDMA-I^Cy5^ with HaCat cell membranes

HaCat cells were plated onto glass-bottom dishes at a density of 1×10 ^5^ cells per dish and incubated for 24 h. The HaCat cells were incubated with OPDMA-I**^Cy5^** (Cy5-equivalent concentration at 0.1 μg/mL) for 6 h, 12 h, and 24 h. According to the experimental needs, the cell membrane was stained with NBD-C6-HPC (1 μM) for 5 min, and then the cells were washed with PBS three times and observed under CLSM. For time-lapse videos, the cells were imaged immediately after adding OPDMA-I^Cy5^ (Cy5-eq. concentration at 0.1 μg/mL). Fluorescent images were taken using CLSM with excitation at 488 nm for NBD-C6-HPC and 640 nm for Cy5.

### Observation of OPDMA-I ^Cy5^ direct transfer between HaCat cells

HaCat cells were seeded in 6-well plates at 1.25 or 2.5× 10^5^ cells per well and allowed to adhere overnight. The cells were then incubated with OPDMA-I^Cy5^ (Cy5-equivalent concentration at 0.1 μg/mL) for 24 h. The OPDMA-I^Cy5^ treated-HaCat cells were then extensively rinsed with sterilized PBS and plated with untreated HaCat-GFP cells. The cells were co-cultured further 24 h and imaged with CLSM with excitation at 488 nm for GFP and 640 nm for Cy5 and analyzed by flow cytometry; the transfer efficacy of OPDMA-I^Cy5^ between cells was calculated according to cell counts.

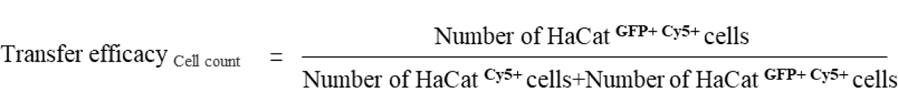

## Supplementary Figures

**Scheme 1.**
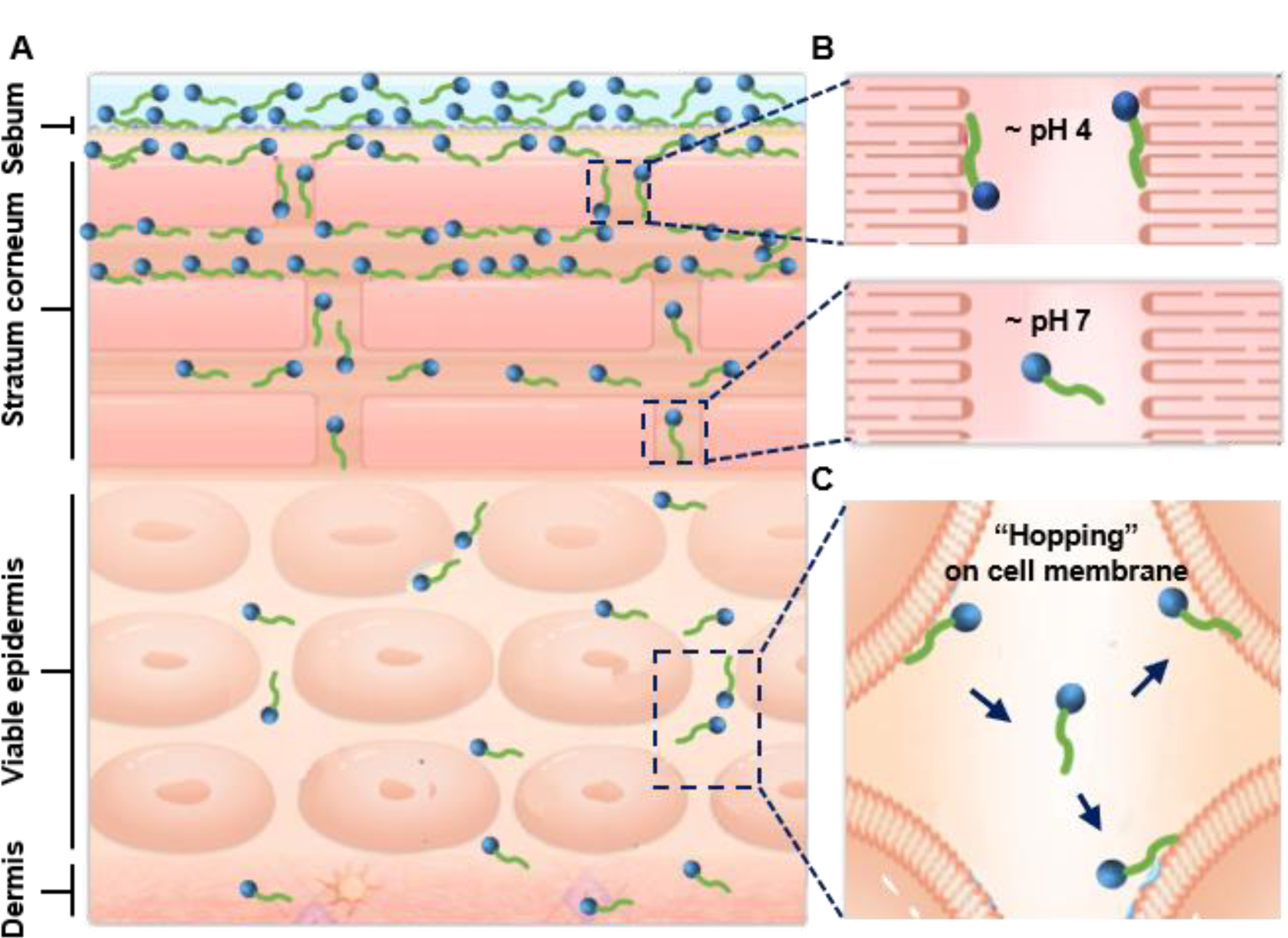
Schematic illustration of the skin penetration after transdermal application of OPDMA-I. **A,** OPDMA-I primarily penetrated the skin epidermis barrier via the paracellular pathway. **B,** OPDMA-I exhibited binding and partitioning abilities with highly structured intercellular lipid lamellae due to the electrostatic effect, leading to efficient diffusion through the SC. **C,** OPDMA-I could traverse the viable epidermis by cell membrane-dependent direct transfer.

**Fig. S1.**
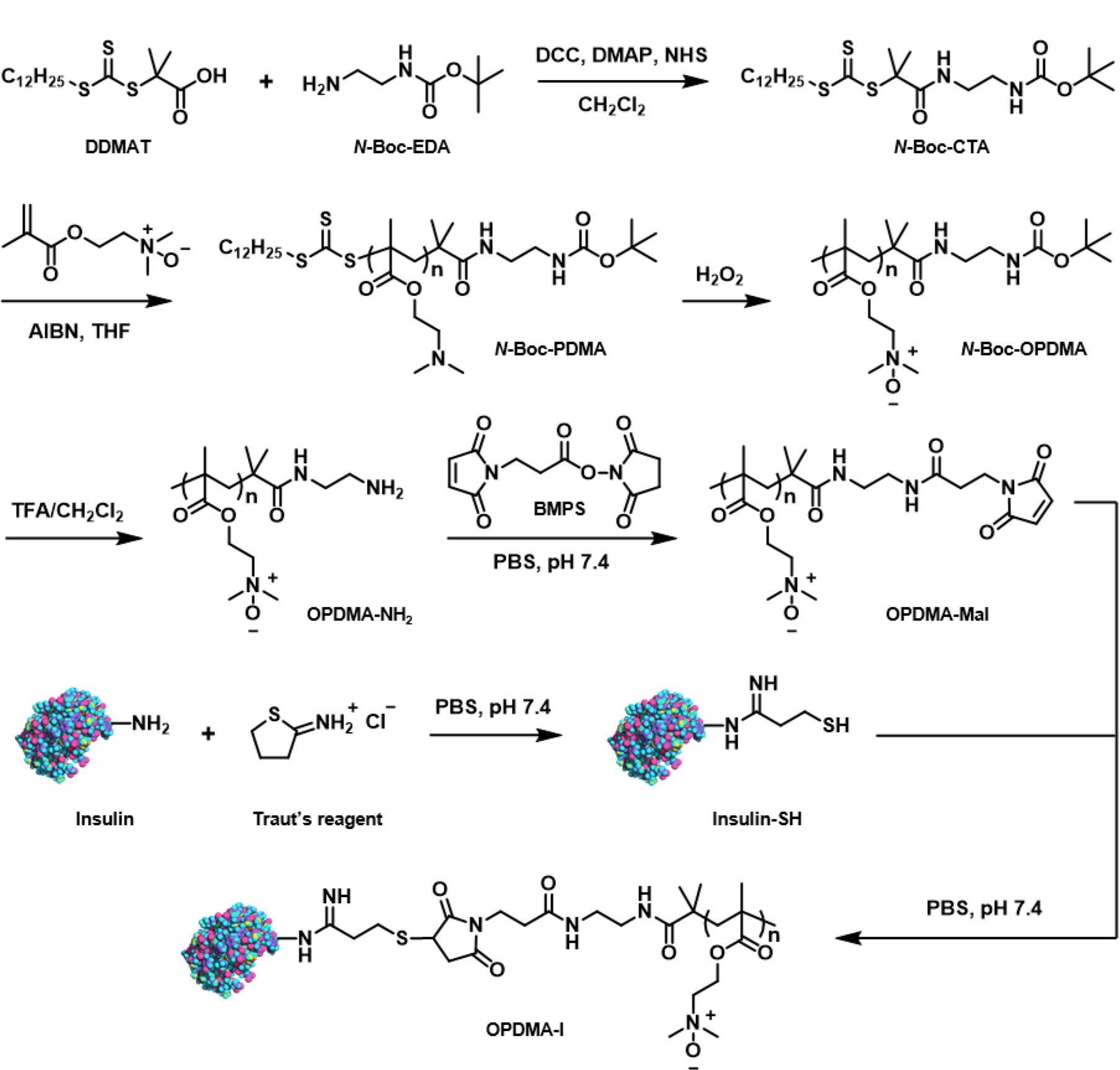
The synthetic scheme of OPDMA-I.

**Fig. S2.**
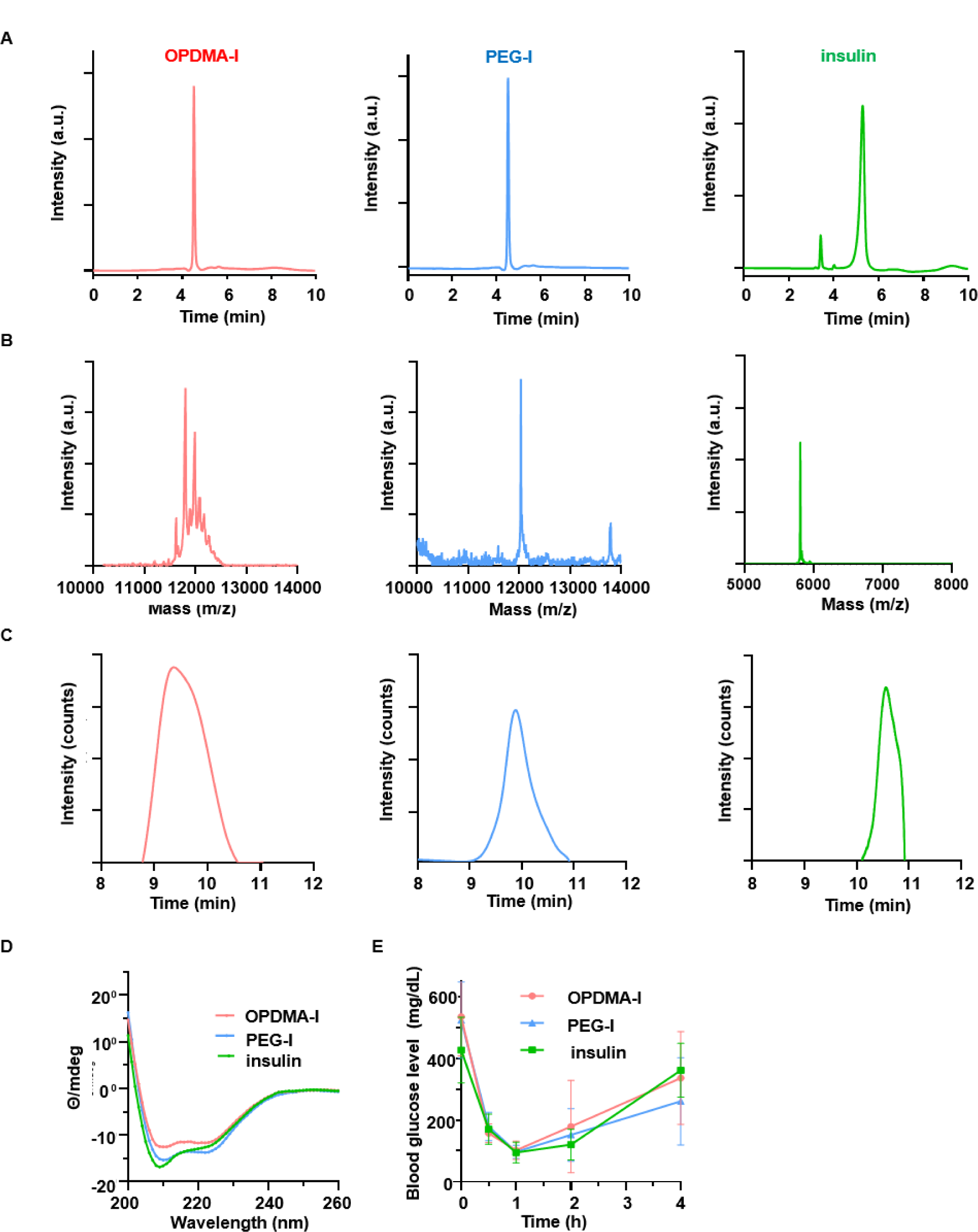
Characterization of OPDMA-I. **A,** Reverse phase HPLC chromatograms with absorbance at 214 nm of OPDMA-I, PEG-I, and native insulin (I). **B,** MALDI-TOF MS spectra of OPDMA-I, PEG-I, and native insulin (I). **C,** Gel permeation chromatography (GPC) traces of OPDMA-I, PEG-I, and native insulin (I) in H_2_O. **D**, Circular dichroism (CD) spectra of native insulin, OPDMA-I, and PEG-I at pH 7.4. **E,** Blood glucose levels of s.c. administrated OPDMA-I, PEG-I, and native insulin in diabetic mice. Insulin-eq. dose: 5U/kg. Data are presented as mean ± SD. (n= 5).

**Fig. S3.**
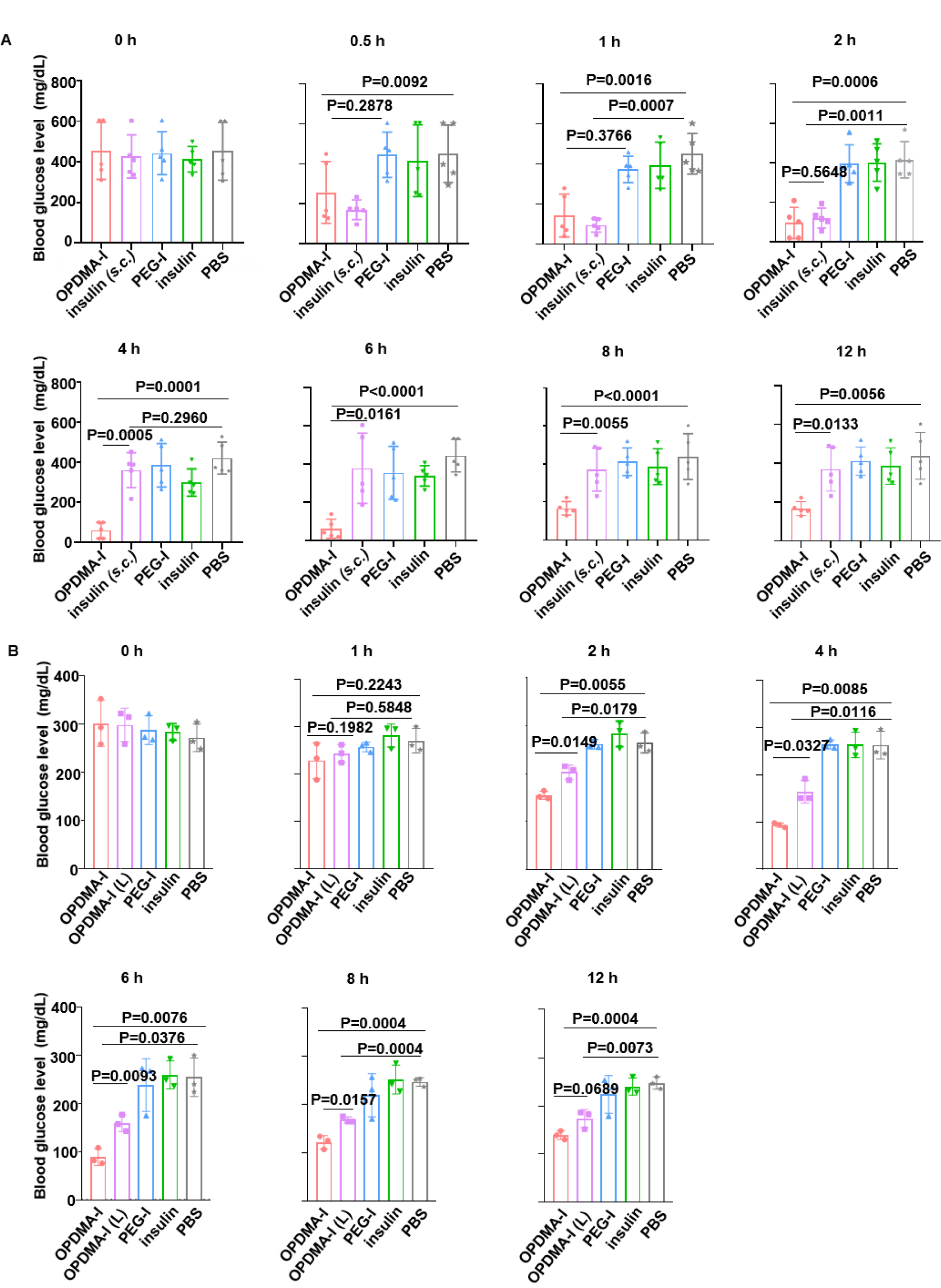
**A,** Individual blood glucose level of each time point in Fig.1B (n= 5). **B**, Individual blood glucose level of each time point in Fig.1H (n= 5).

**Fig. S4.**
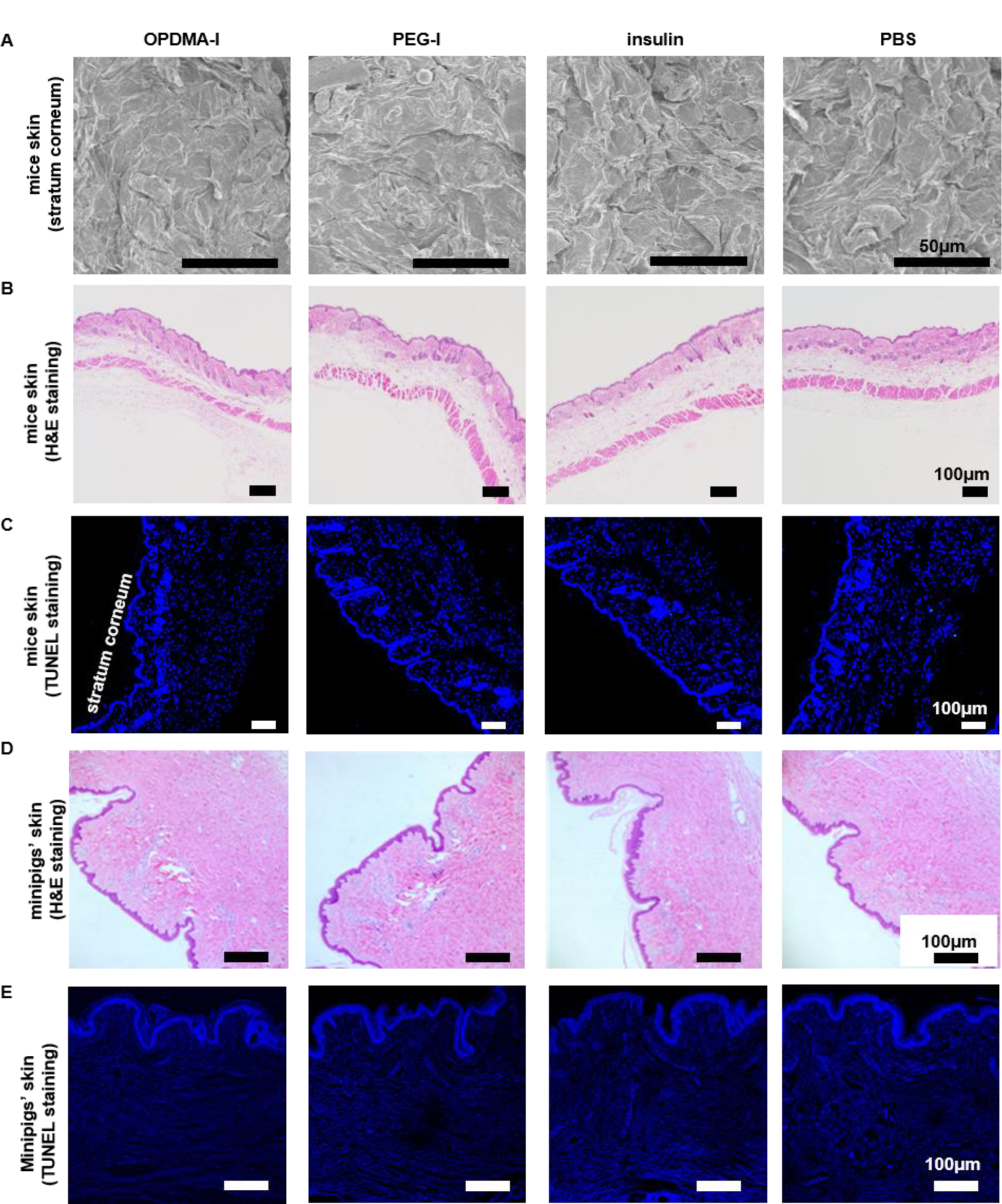
**A**, SEM images of mice skin surface after treatment with OPDMA-I, PEG-I, native insulin, or PBS. Scale bars, 50 μm. **B**, Representative images of H&E staining results mice skin slices after treatment with OPDMA-I, PEG-I, native insulin, and PBS. Scale bars, 100 μm. **C**, Immunohistology staining with TUNEL assay (green) and Hoechst (blue) of mice skin slices after treatment with OPDMA-I, PEG-I, native insulin, and PBS. Scale bars, 100 μm. **D**, Representative images of H&E staining results in the treated site of minipig skin. Scale bars, 100 μm. **E**, Immunohistology staining with TUNEL assay (green) and Hoechst (blue) of results in the treated site of minipig skin. Scale bars,100μm.

**Fig. S5.**
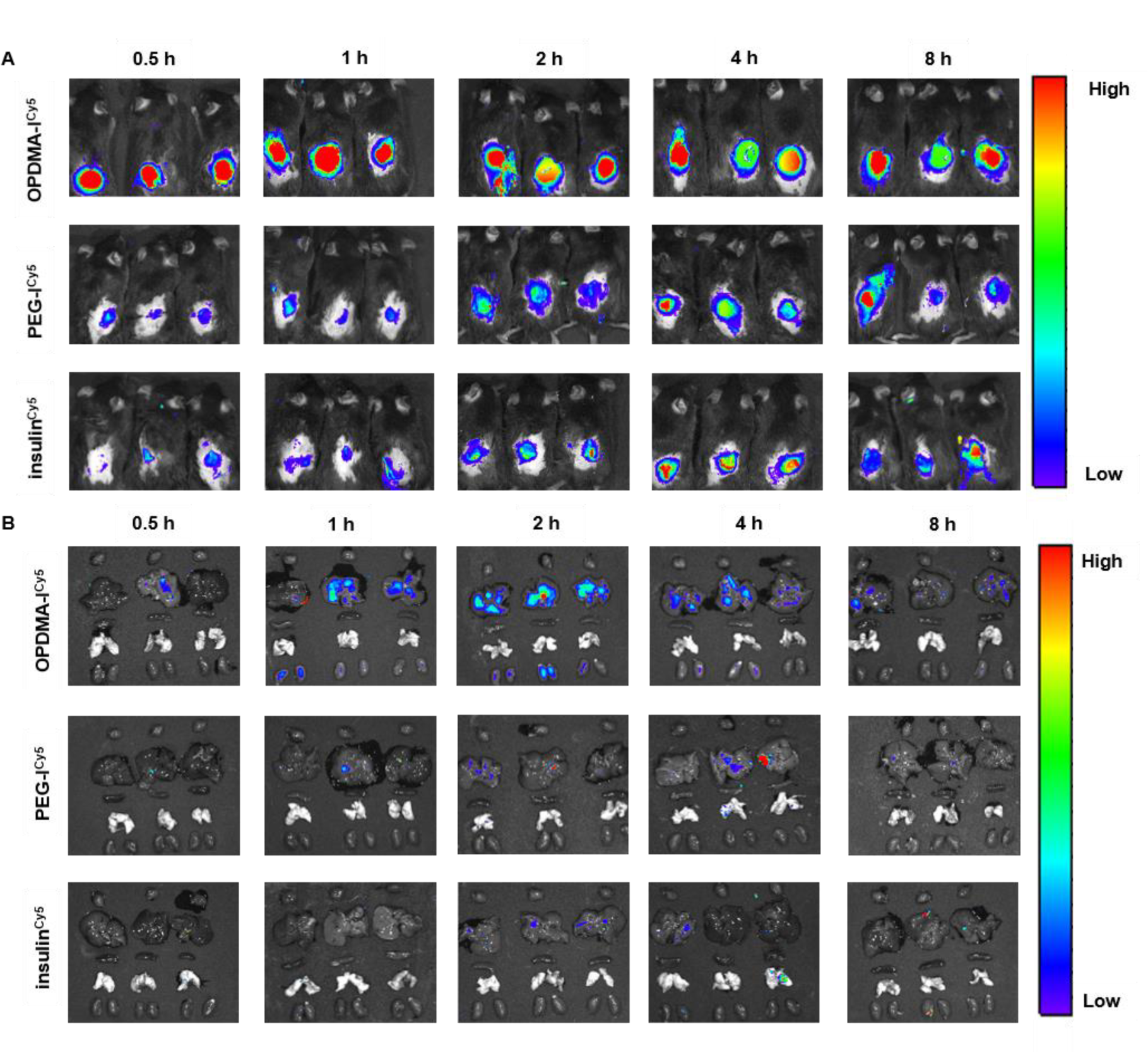
**A,** Fluorescence images of mice after transdermal administration of OPDMA-I^Cy5^, PEG-I^Cy5^, and insulin^Cy5^ on dorsal skin ( Cy5-eq. dose: 1μg/mL, application area: 1.77 cm^2^). **B,** Fluorescence images of mice tissue after transdermal administration of 0.2 mL OPDMA-I^Cy5^, PEG-I^Cy5,^ and insulin^Cy5^ solutions (Cy5-eq. concentration: 1μg/mL) at the dorsal skin with an area of 1.77 cm^2^.

**Fig. S6.**
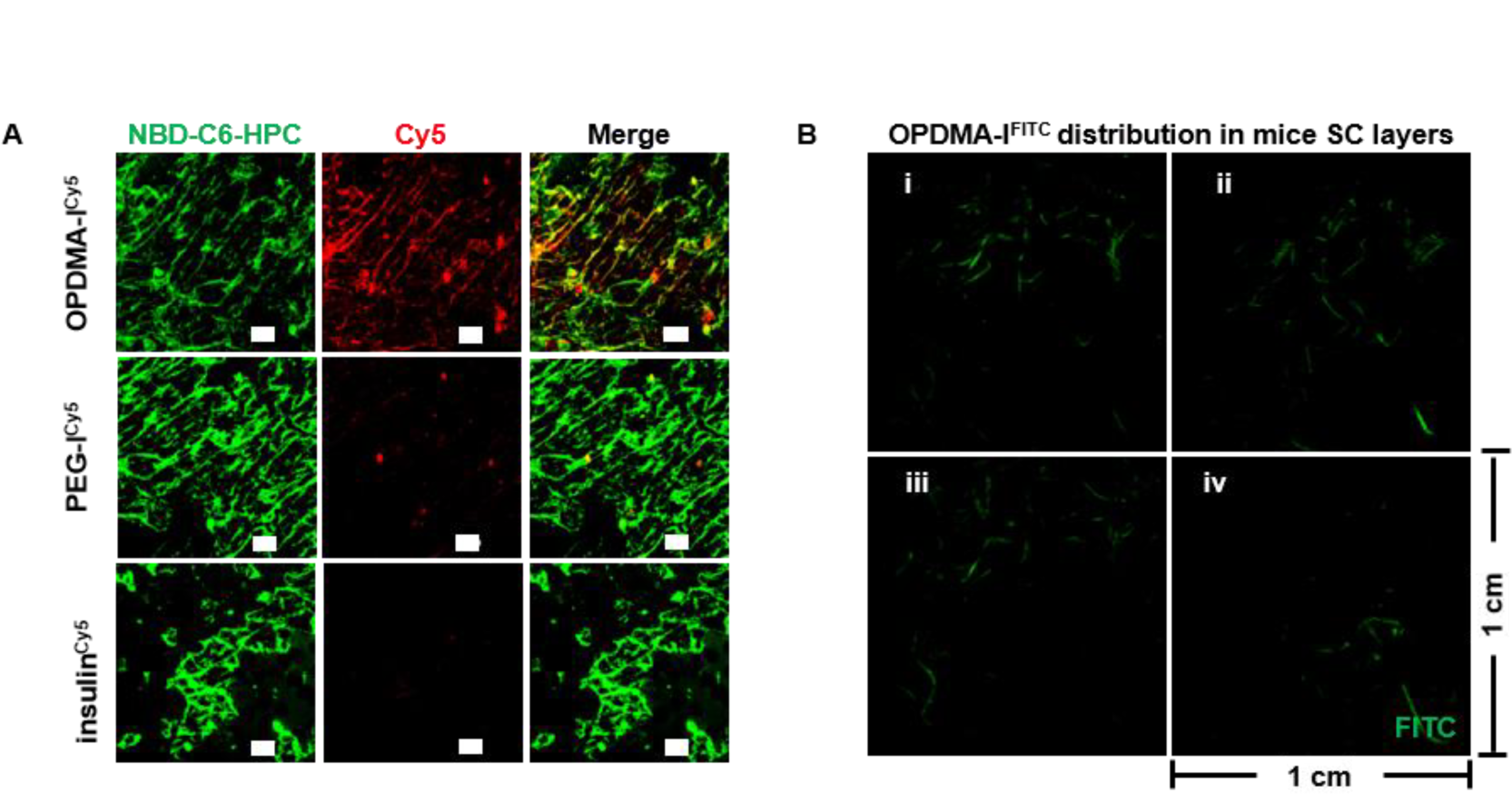
**A,** Fluorescence distribution of insulin^Cy5^, PEG-I^Cy5^, or OPDMA-I^Cy5^ (red) in NBD-C6-HPC stained SC intercellular lipid (green) imaged by CLSM. SC samples were extracted from the deep layer of mice’s dorsal skin by adhesive tape after in vivo skin permeation for 4 h-post transdermal application of 0.2 mL insulin solution. (Cy5-eq. concentration, 1 μg/mL, application area: 1.77cm^2^). Scale bars, 25 μm. **B,** Representative sequential images of 0.2 mL OPDMA-I^FITC^ treated mice SC layers. (FITC-eq. dose: 1μg/mL, application area: 1.77 cm^2^).

**Fig. S7.**
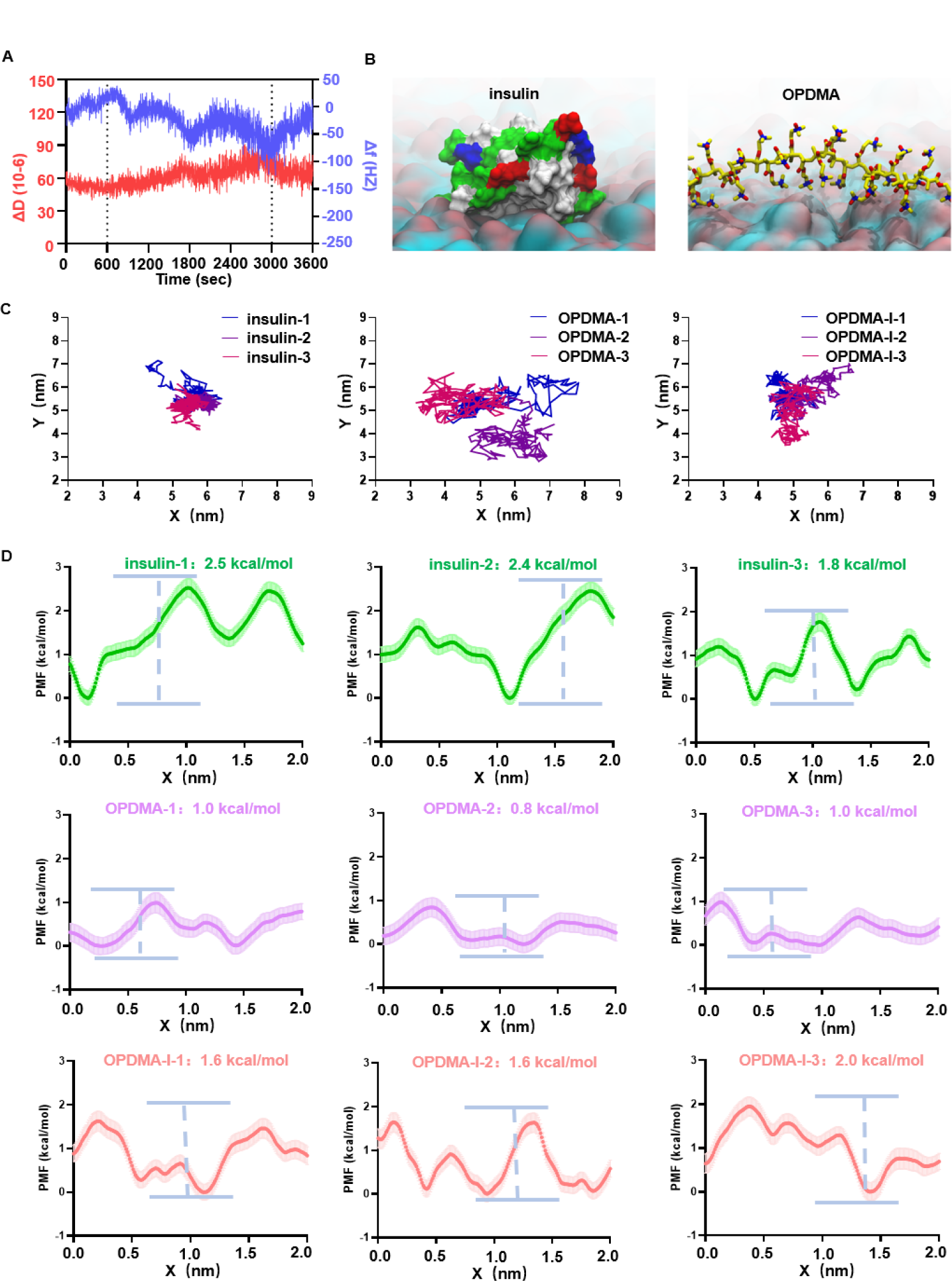
**A,** Changes in resonant frequency (F) and dissipation (D) of model SC lipid membrane-coated quartz crystal of the attachment of OPDMA-I solution (insulin-eq. concentration:0.5 mg/mL). **B,** Representative local binding modes of insulin and OPDMA on the rough SC lipid surfaces. It was shown that insulin could be anchored on the inhomogeneous lipid surface with a preferential binding mode. **C,** Traces of insulin, OPDMA, and OPDMA-I on SC lipids in three 100-ns independent molecular dynamics trajectories for each of them. **D,** PMF analysis along the surface of SC lipids in different paths to estimate the energy barriers for diffusion, which were 2.2 ± 0.3 kcal/mol for insulin, 0.9 ± 0.1 kcal/ mol for OPDMA and 1.7 ± 0.2 kcal/mol for OPDMA-I.

**Fig. S8.**
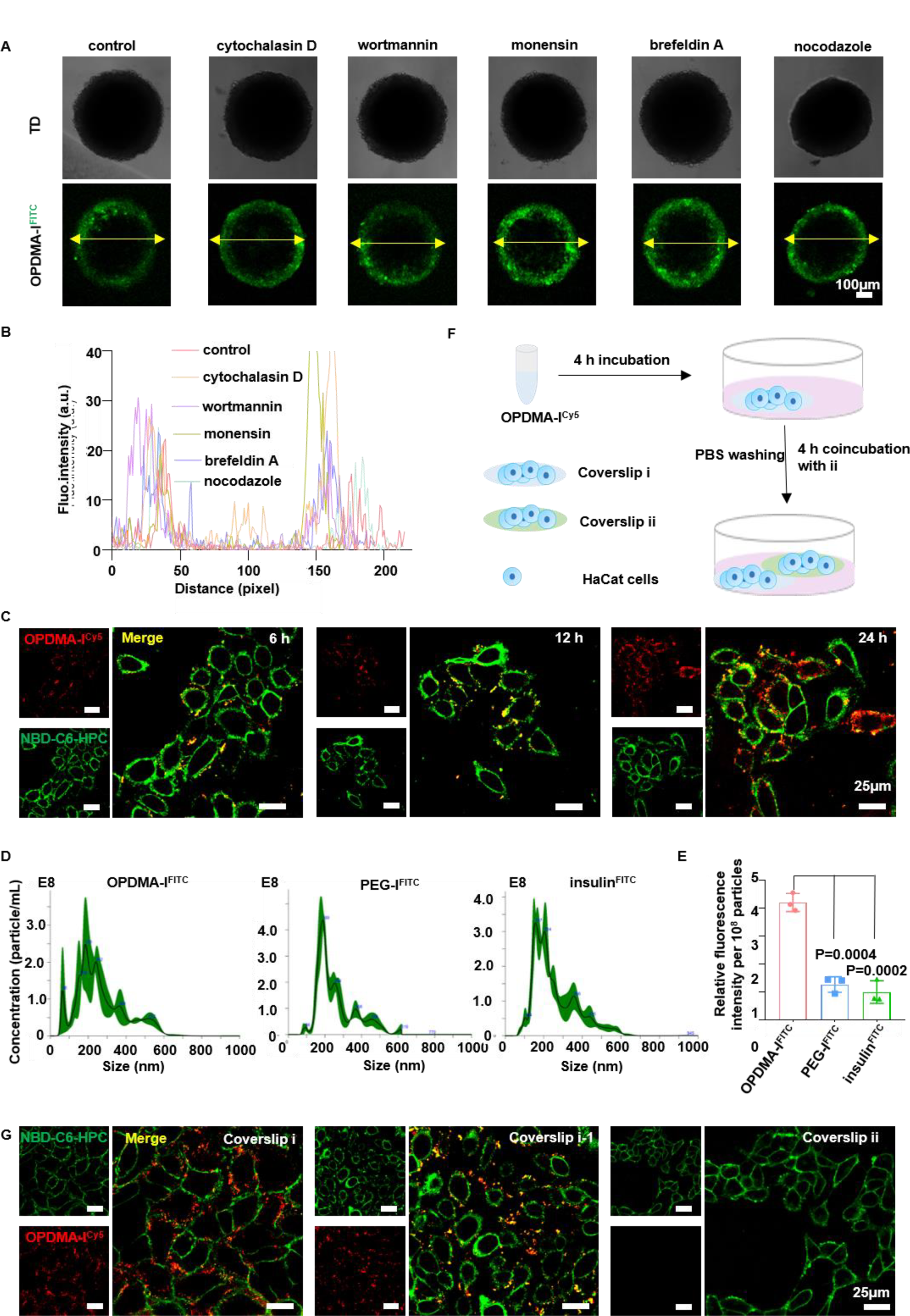
**A-B,** HaCat spheroids were separately pre-treated with the inhibitors for 4 h and OPDMA-I^FITC^ (FITC-eq. dose, 0.1 μg/mL) for 4 h, and then imaged with CLSM by Z-stack tomoscan at 20 μm intervals. **A,** Representative images of OPDMA-I^FITC^ distribution in the spheroids, five images obtained per group. Scale bars,100 μm. **B,** FITC fluorescence intensity along the yellow arrows in **A**. **C,** Cell membrane absorption of OPDMA-I^Cy5^ on NBD-C6-HPC stained HaCat cells observed by CLSM. Representative images at 6 h,12 h, and 24 h are shown. Scale bars, 25 μm. Cy5-eq. dose, 0.1 μg/mL. **D,** The particle size distribution of OPDMA-I^FITC^, PEG-I^FITC,^ and insulin^FITC^ treated HaCat-Derived Nanovesicles measured by nanoparticle tracking analysis (NTA); HaCat cells were incubated with OPDMA-I^FITC^, PEG-I^FITC^, and insulin^FITC^ for 24 h with FITC-eq. dose of 0.1 μg/mL. **E,** Representative relative FITC fluorescence intensity of HaCat-Derived Nanovesicles in e detected by a microplate reader. Data are mean ± SD (n= 3). **F-G,** Transcellular transfer of OPDMA-I^Cy5^ visualized by CLSM. HaCat cells (∼10^5^) on coverslip (i) were cultured in a medium that contained OPDMA-I^Cy5^ ( Cy5-eq. dose: 0.1 μg/mL) for 4 h. Coverslip (i) was rinsed and imaged and then put into fresh culture medium along with coverslip (ii) with fresh cells for 12 h. The cell membrane was stained with NBD-C6-HPC. Scale bars, 25 μm.

**Table S1.** Serum ALB, Tp, GLO, AST, and ALP concentration and RBC, HGB, MCV, MCHC, and RDW were counted after the whole treatment of minipigs. Data are presented as mean ± SD (n = 3).

| Biochemical indicators |  | Blood routine |  |
| --- | --- | --- | --- |
| Albumin (ALB) (g/L) | 36.3±0.3 | Red blood cells (RBC) ($10^{12}$/L) | 6.6±1.0 |
| Total protein (Tp) (g/L) | 73.2±0.2 | Hemoglobin (HGB) (g/L) | 122.6±1.4 |
| Globulin (GLO) (g/L) | 36.0±1.0 | Mean red blood cell volume (MCV) (fL) | 54.4±1.4 |
| Serum aspartate aminotransferase (AST) (U/L) | 20.3±2.1 | Mean red blood cell hemoglobin concentration MCHC(g/L) | 347.3±4.0 |
| Serum alkaline phosphatase (ALP) (U/L) | 212.0±1.0 | Red blood cell distribution width (RDW) (%) | 19.9±1.9 |

**Table S2.** Summary of penetration parameters of OPDMA-I across EpiKutis® model. Data are shown with mean ± SD (n=3).

| | The cumulative amount of permeation in 24 h (μg) | $J_{ss}$ (μg/cm<sup>2</sup>/h) | $K_p$ (cm/h) | Enhancement ratio of $K_p$ |
| --- | --- | --- | --- | --- |
| OPDMA-I | 14.52±2.41 | 0.50±0.12 | 2.94$\times 10^{-3}$ | 9.17 |
| PEG-I | 2.45±0.05 | 0.10±0.01 | 0.6$\times 10^{-3}$ | 2.04 |
| insulin | 1.47±0.12 | 0.05±0.01 | 0.3$\times 10^{-3}$ | 1.00 |

